# An unusual nucleosomal sequence pattern is enriched in mammalian genes

**DOI:** 10.1101/207167

**Authors:** Gregory M. Wright, Feng Cui

## Abstract

Nucleosomal DNA sequences generally follow a well-known pattern with ∼10-bp periodic WW (where W is A or T) dinucleotides that oscillate in phase with each other and out of phase with SS (where S is G or C) dinucleotides. However, nucleosomes having a DNA sequence pattern inverse to the conventional one have not been systematically analyzed in eukaryotes. Here, we show that anti-WW/SS nucleosomes are widespread and exhibit a species- and context-specific distribution in the genomes. The enrichment of anti-WW/SS nucleosomes in mammalian genomes is positively correlated with RNA Pol II transcriptional levels, but negatively correlated with the presence of the periodic WW or SS sequence patterns. In addition, chromatin remodeling complexes have an impact on the abundance of anti-WW/SS nucleosomes. The data reveal distinct roles of *cis*- and *trans*-acting factors in the rotational positioning of nucleosomes between non-mammals and mammals. Implications of the anti-WW/SS sequence pattern for RNA Pol II transcription are discussed.

## Introduction

The eukaryotic genome is organized into chromatin in which the access of protein to DNA is tightly regulated to facilitate genomic functions such as transcription, replication and repair. The basic repeating unit of chromatin is the nucleosome core particle (NCP) that consists of a histone octamer, around which 147 base pairs (bp) of DNA is wrapped in ∼1.7 turns (Luger et al. 1997). One of the most intriguing questions in chromosome biology is to understand where nucleosomes are positioned and how specific positioning of nucleosomes affects transcription factor (TF) binding and gene expression in the context of chromatin. Experimentally, nucleosome positions are usually mapped by two types of methods. The first type involves treating chromatin with micrococcal nuclease (MNase) followed by massively parallel sequencing (denoted as MNase-Seq). However, MNase has strong sequence preferences: it cuts predominantly within AT-rich sequences in both free DNA (Dingwall and Lomonossoff 1981; Horz and Altenburger 1981) and in the linker DNA between nucleosomes (Field et al. 2008; Cui and Zhurkin 2009). This sequence specificity can be overcome by a method based on site-directed hydroxyl radicals (denoted as chemical method) (Flaus et al. 1996). The chemical method allows the mapping of nucleosome dyad positions at a single base-pair resolution (Panetta et al. 1998; Flaus and Richmond 1998; Kassaboy et al. 2002) and recently was applied to the yeast (Brogaard et al. 2012; Moyle-Heyrman et al. 2013; Chereji et al 2018) and mouse (Voong et al. 2016) genomes.

Historically, nucleosome positioning is usually characterized by two parameters: rotational positioning, defined by the side of the DNA helix that faces the histones, and translational positioning, defined by the nucleosome midpoint (or dyad) with regard to the DNA sequence (Lu et al. 1994). Several *cis* and *trans* determinants of the translational positioning of nucleosomes have been reviewed in literature (Struhl and Segal 2013).

With regard to the rotational positioning, it is well established that DNA sequence is a critical determinant (Trifonov 2011) and various sequence patterns have been proposed (Mengeritsky and Trifonov 1983; Zhurkin 1983; Drew and Travers 1985; Calladine and Drew 1986; Uberbacher et al. 1988; Ioshikhes et al. 1992; Baldi et al. 1996; Ioshikhes et al. 1996; Lowary and Widom 1998; Kogan et al. 2006). One of the most well-known patterns is the WW/SS pattern (where W is A or T and S is G or C), which was first described by Travers and colleagues (Satchwell et al. 1986). Specifically, WW dinucleotides tend to occur at the sites where nucleosomal DNA bends into minor grooves (*i.e.*, minor-groove bending sites or minor-GBS) facing toward the histone core, while SS dinucleotides are often positioned at the sites where nucleosomal DNA bends into major grooves (*i.e.*, major-groove bending sites or major-GBS) facing toward the histone core. This pattern has been successfully used to predict the rotational positioning of nucleosomes (Cui et al. 2011). An anti-WW/SS pattern was described in which WW runs inverse to SS, and it was found that promoter nucleosomes in yeast favor this pattern (Ioshikhes et al. 2011). That is, the number of nucleosome following the anti-WW/SS pattern exceeds the number of nucleosomes following the conventional WW/SS pattern in yeast promoters.

Structurally, highly conserved “sprocket” arginine residues (Muthurajan et al. 2003; Sullivan and Landsman 2003) insert into the minor-GBS (Davey et al. 2002). Short poly(dA:dT) stretches help to narrow DNA minor grooves, potentially enhancing electrostatic interactions between DNA phosphate backbone and “sprocket” arginine residues (Rohs et al. 2009; Wang et al. 2010; West et al. 2010). These arginine-DNA contacts that occur every ∼10 bp within NCP provides the structural basis for the rotational positioning of nucleosomes (Luger et al. 1997; Rohs et al. 2009; Wang et al. 2010; West et al. 2010). In this light, anti-WW/SS nucleosomes seem to have unfavorable DNA-histone interactions, representing a relatively unstable structure. However, many questions of how anti-WW/SS nucleosomes are distributed across eukaryotic genomes and what factors influence their abundance remain poorly understood.

Here, we have quantified nucleosomes with or without the WW/SS sequence pattern, and systemically analyzed the fraction and the distribution of anti-WW/SS nucleosomes across five eukaryotic genomes under different cellular and growth conditions. We found that anti-WW/SS nucleosomes are widespread in the genomes regardless of the mapping methods and distributed in a species- and context-specific manner. In non-mammals including yeast, anti-WW/SS nucleosomes are not enriched in promoters. However, in mammals, this type of nucleosomes is enriched in promoter and genic regions but not in repetitive DNA elements. The enrichment of anti-WW/SS nucleosomes is positively correlated with RNA Pol II transcriptional levels, but negatively correlated with the presence of the periodic WW (or SS) pattern. Moreover, we found that chromatin remodelers have an impact on the number but not the distribution of anti-WW/SS nucleosomes in non-mammals. Our results indicate that *cis-* and *trans*-acting factors play distinct roles in controlling the rotational positioning of nucleosomes between non-mammals and mammals.

## Results

### Anti-WW/SS nucleosomes are widespread in eukaryotes

NCP fragments of 147 bp in length were divided into 4 sequence patterns, Type 1-4 (Table 1), based on the relative occurrence of WW and SS dinucleotides in 12 minor- and 12 major-GBS (Supplementary Table S1, Supplementary Fig. S1). The Type 1 pattern represents the conventional WW/SS sequence pattern in which WW and SS preferentially occur in minor-GBS and major-GBS respectively (Fig. 1A). Type 2 nucleosomes follow a ‘mixed’ pattern, in which both WW and SS dinucleotides are more abundant in minor-GBS than major-GBS (Fig. 1B). The Type 3 pattern is opposite to the Type 2 pattern in which both WW and SS are more abundant in major-GBS than minor-GBS (Fig. 1C). The Type 4 pattern is inverse to Type 1 with WW and SS dinucleotides preferentially occurring in major- and minor-GBS respectively (Fig. 1D). For the sake of simplicity, nucleosomes with the Type 1 pattern are denoted as WW/SS nucleosomes, whereas those with the Type 4 pattern are denoted as anti-WW/SS nucleosomes.

**Table 1.** Classification of nucleosomal DNA sequence patterns.

| Sequence Type | WW | SS |
| --- | --- | --- |
| Type 1 | **minor** WW ≥ **Major** WW * coef<sup>†</sup> | **minor** SS ≤ **Major** SS * coef |
| Type 2 | **minor** WW ≥ **Major** WW * coef | **minor** SS $>$ **Major** SS * coef |
| Type 3 | **minor** WW $<$ **Major** WW * coef | **minor** SS ≤ **Major** SS * coef |
| Type 4 | **minor** WW $<$ **Major** WW * coef | **minor** SS $>$ **Major** SS * coef |

<sup>†</sup>coef: the coefficient is a constant of 1.125 (=36/32). It refers to the ratio between 36 dinucleotide positions in minor-groove bending sites and 32 dinucleotide positions in major-groove bending sites (see Supplemental Table S1).

**Figure 1.**
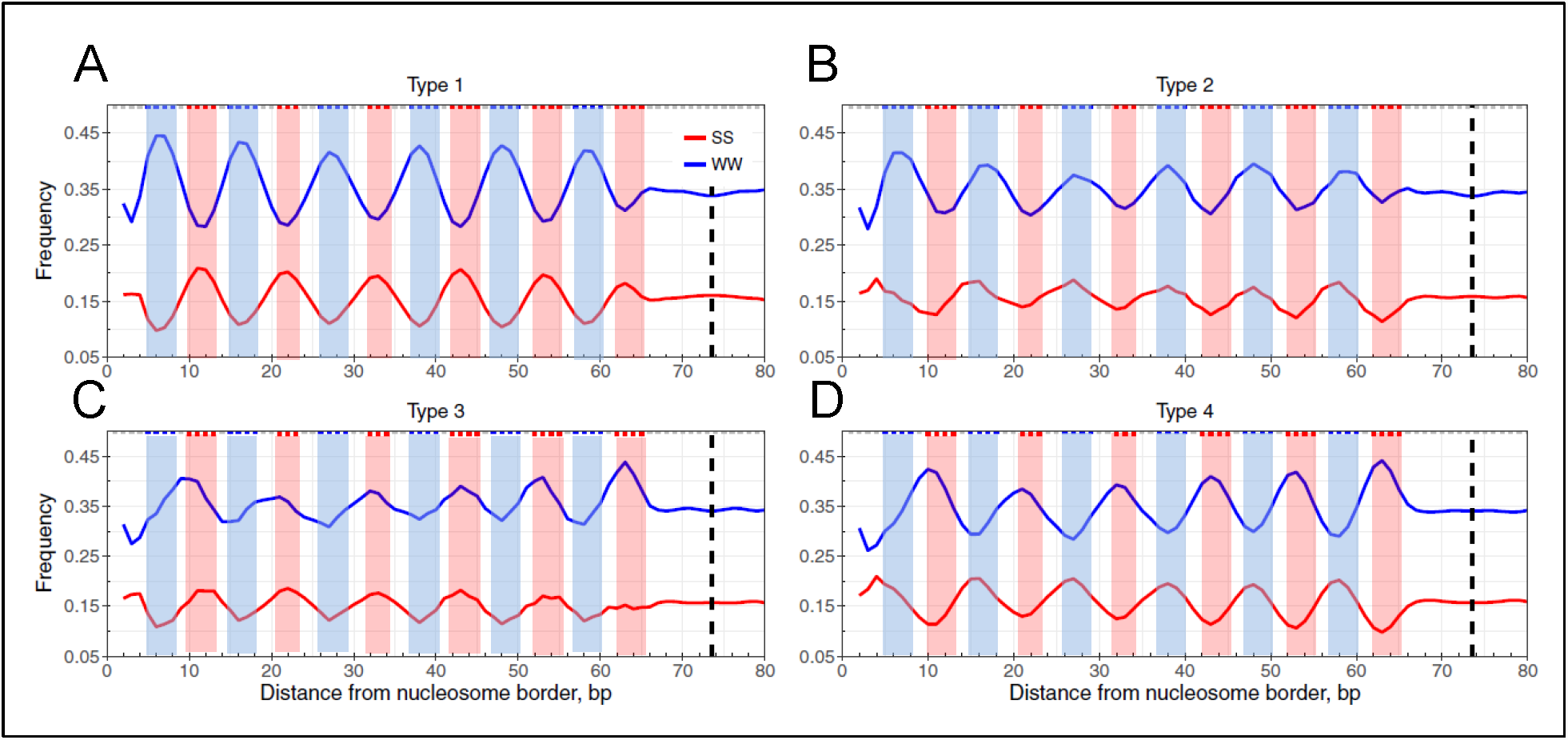
Four sequence patterns of nucleosomal DNA in yeast. Shown are frequencies of the combined AA, TT, AT, and TA dinucleotides (denoted as WW, shown in blue) and GG, CC, GC and CG dinucleotides (denoted as SS, shown in red) at each nucleosomal position, which were ‘symmetrized’ with respect to the dyad (dashed lines). Three base-pair running averages of the WW and SS frequencies were calculated and plotted: Type 1 (A), Type 2 (B), Type 3 (C) and Type 4 (D). The blue and red shade areas cover the range of minor- and major-GBS, respectively (see Methods).

We have identified all four types of sequence patterns in datasets (Supplementary Tables S2-S3) from yeast, fruit flies, nematodes, mice and humans (Fig. 1 and Supplementary Fig. 2-7). Our analysis has led to several interesting observations. First, the datasets were produced by either the MNase-Seq method or the chemical method, indicating that the presence of anti-WW/SS nucleosomes is independent of nucleosome mapping methods. Second, anti-WW/SS nucleosomes occur in all chromosomes (Supplementary Table S4), indicating that they are widespread in the genomes. Third, except for the yeast dataset obtained by chemical mapping, the fraction of WW/SS nucleosomes is less than 50% of all nucleosomes (Supplementary Fig. 8A), indicating that the canonical WW/SS sequence pattern is not predominant, and the majority of eukaryotic nucleosomes actually follow a non-canonical pattern (Supplementary Fig. 8A). Fourth, the fractions of Type 1 and Type 4 nucleosomes vary substantially. For instance, the fractions of the 4 types of nucleosomes are 36-57% (Type 1), 20-22% (Type 2), 8-13% (Type 3) and 13-31% (Type 4) respectively (Supplementary Table S4). Since Type 2 and Type 3 nucleosomes have relatively small changes over datasets, we were prompted to develop a measure, ΔNPS (= Type 1 (%) – Type 4 (%)), to gauge the difference between Type 1 and Type 4 nucleosomes (Supplementary Fig. 8B). A ΔNPS value represents the relative abundance of WW/SS *versus* anti-WW/SS nucleosomes in a given genomic region.

Taken together, anti-WW/SS nucleosomes accounting for 13-31% of total nucleosomes exist in all eukaryotes examined and are widespread across the genomes. We next sought to understand how anti-WW/SS nucleosomes are distributed in the genomes.

### Anti-WW/SS nucleosomes are enriched in mammalian genes and associated with transcription

We started with genomic regions surrounding the transcriptional start sites (TSS) of genes that were separated into quartiles by transcriptional levels (Supplementary Table S5). As expected, nucleosome occupancy profiles in yeast reveal a well-established genomic pattern (Sekinger et al. 2005; Yuan et al. 2005): the presence of a ∼200-bp nucleosome-depleted region (NDR), flanked by phased nucleosomes, which form a highly regular array extending into gene bodies (Fig. 2A-B). The phased nucleosomes, named as nucleosome −1, +1, +2, +3, +4, and +5, are organized relative to TSS (Mavrich et al. 2008; Shivaswamy et al. 2008) (Supplementary Table S6).

**Figure 2.**
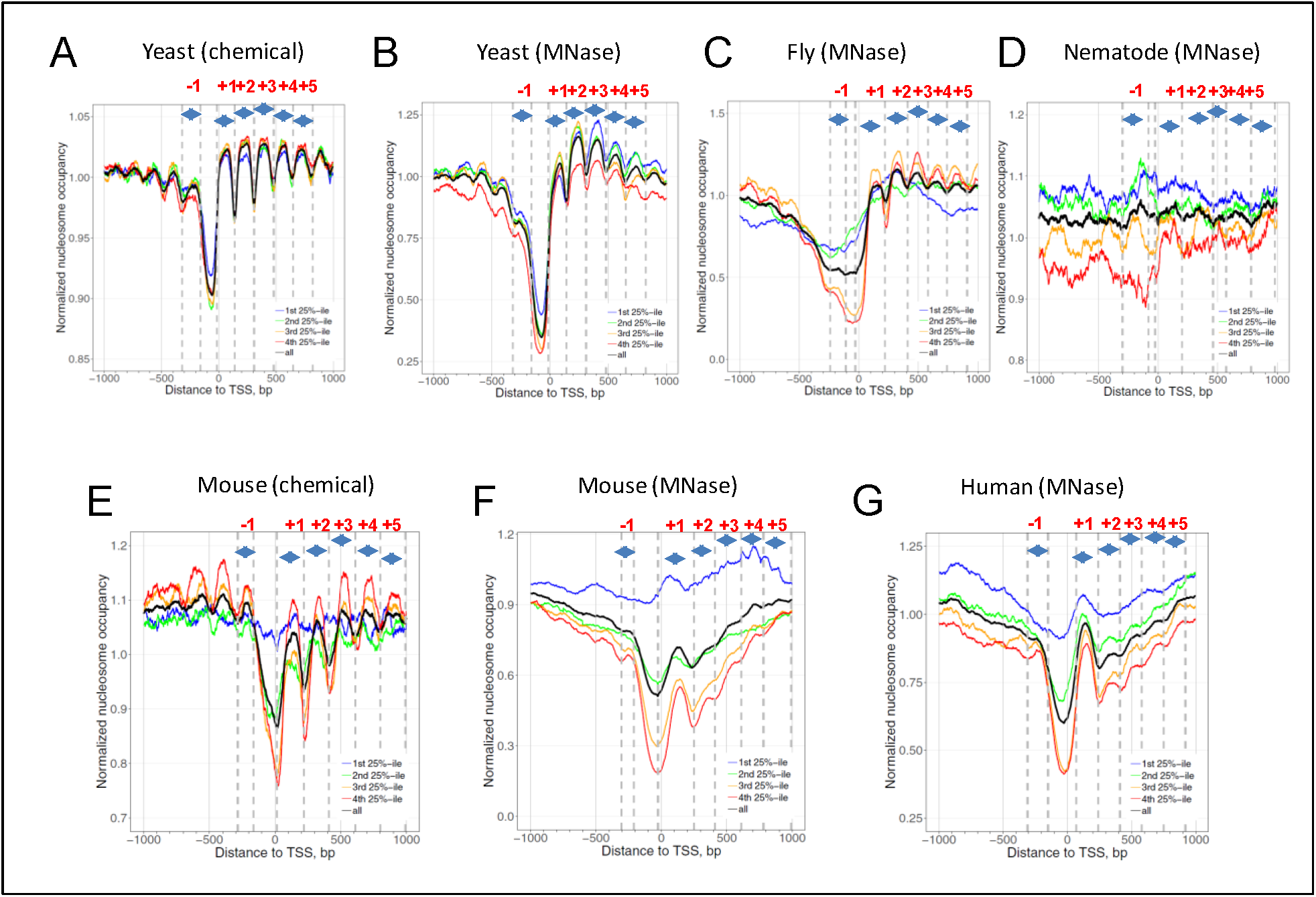
Nucleosome occupancy profiles around TSS of yeast (A-B), fly (B), nematode (C), mouse (D-E) and human (F) genes. There are two separate nucleosome datasets for yeast and mice, derived from chemical mapping (A, D) and MNase-Seq mapping (B, E), respectively. Nucleosome occupancy signals ±1kb of verified TSSs are separated into quartiles based on transcriptional levels (Table S5). The 1^st^ 25%-ile represents the least active genes whereas the 4^th^ 25%-ile represents the most active genes. The average nucleosome profile of all genes is shown in black. Nucleosomes −1 to +5 are demarcated by dashed lines and arrows, following the methods used in previous studies (Jiang and Pugh 2009). These definable zones relative to the TSS (position 0) to which a nucleosome midpoint may be assigned are: −1, +1, +2, +3, +4, and +5 (see Table S6 for nucleosome ranges).

Overall, the regularity of the nucleosome arrays depends on species, transcription levels, and nucleosome mapping methods (Fig. 2). First, nucleosome arrays in yeast are more regular than their counterparts in other species. Second, as transcriptional frequencies go higher, the nucleosome arrays tend to be more regular (compare red lines with blue lines). Third, nucleosomes mapped by the chemical method tend to more regularly positioned than those mapped by the MNase-Seq method (compare Fig. 2A-B with Fig.2E-F), presumably because the hydroxyl-cleavage-based chemical method overcomes the sequence specificity of MNase (Dingwall and Lomonossoff 1981; Horz and Altenburger 1981) and is able to map nucleosomes at a higher resolution (Flaus et al. 1996).

Comparison of nucleosomal ΔNPS value for all genes across the species reveals some interesting patterns. First, in the non-mammalian genes, the ΔNPS values appear similar to their genomic ΔNPS (see black lines and dashed lines in Fig. 3A-D), suggesting that the relative abundance of Type 1 and Type 4 nucleosomes has little change. This pattern is confirmed by additional datasets in flies (Supplementary Fig. 9) and nematodes (Supplementary Fig. 10). Consistently, quantitative analysis reveals small difference between nucleosomal ΔNPS (averaged over nucleosome −1 to +5) and genomic ΔNPS: < 1% for yeast and 1-3% for flies and nematodes (Supplementary Table S7). Apparently, our data do not support previous observations that anti-WW/SS nucleosomes are enriched in yeast promoters, and the number of anti-WW/SS nucleosomes exceeds that of WW/SS nucleosomes (Ioshikhes et al. 2011).

**Figure 3.**
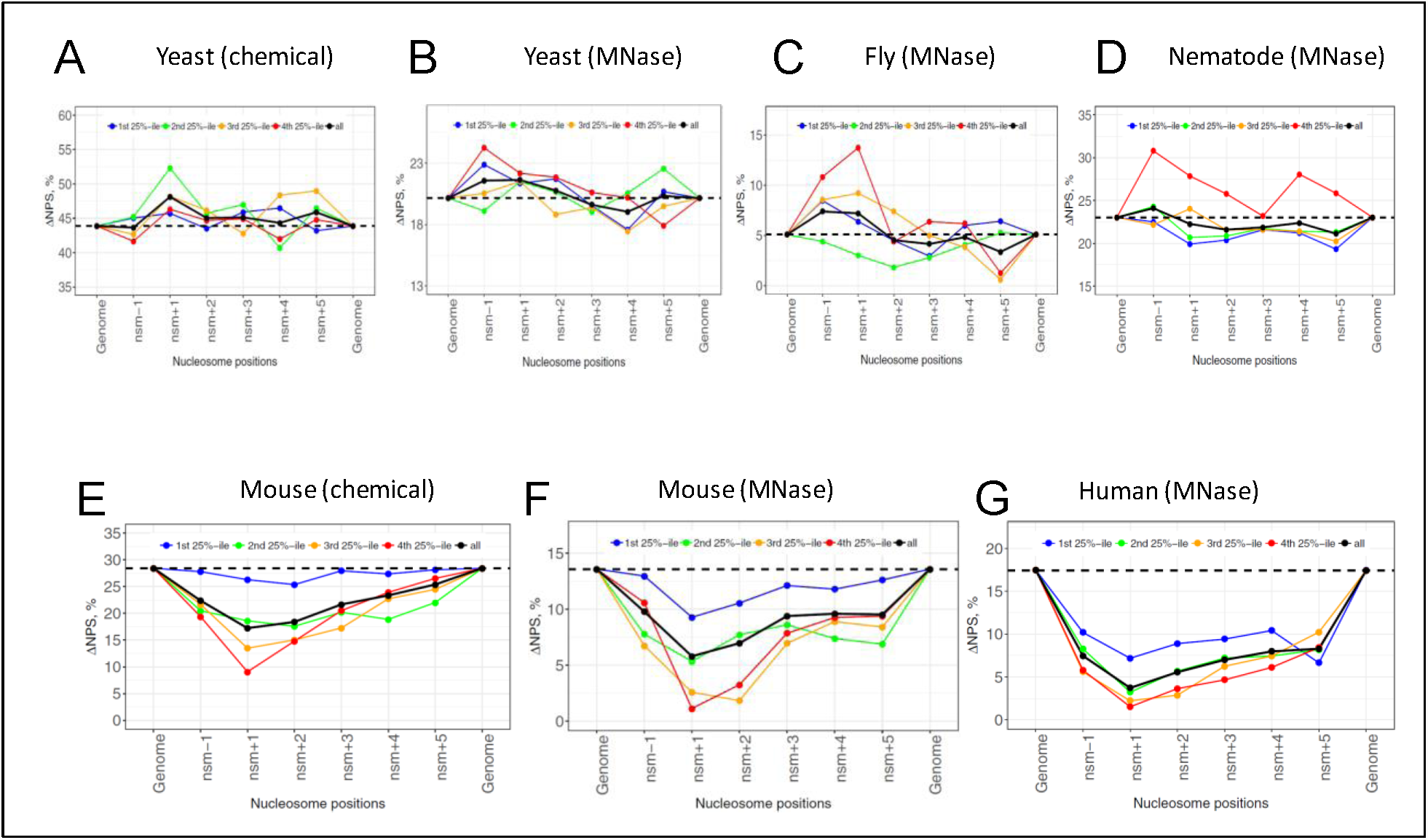
Nucleosome ΔNPS values for genes separated into quartiles by transcriptional frequencies. The 1^st^ 25%-ile represents the least active genes whereas the 4^th^ 25%-ile represents the most active genes. The ΔNPS values for all genes are shown in black. The genomic ΔNPS values are denoted by dashed lines.

Second, in mammalian genes, the ΔNPS values of nucleosome −1 to +5 tend to be lower than the genomic ΔNPS, with the lowest point at nucleosome +1 (black lines in Fig. 3E-G), indicating that the relative abundance of Type 1 and Type 4 nucleosomes have changed. The averaged nucleosomal ΔNPS values exhibit a substantial difference, 5-11%, from the genomic ΔNPS value, which is much higher than that of non-mammals, 0-3% (Supplementary Table S7). Detailed analysis on the ΔNPS values of nucleosome +1 in the mouse chemical dataset shows that the fraction of Type 1 nucleosomes is decreased from 47% to 41%, whereas the fraction of Type 4 nucleosomes is increased from 19% to 24% (Supplementary Table S8). A similar trend is seen in the mouse and human MNase datasets (Supplementary Table S8), clearly illustrating that the fraction of Type 1 nucleosomes is decreased whereas the fraction of Type 4 nucleosome is increased in nucleosome +1. This mammalian pattern (Fig. 3E-G) is in marked contrast to the non-mammalian one (Fig. 3A-D), showing that anti-WW/SS nucleosomes are enriched in mammalian genes.

Third, the change in ΔNPS values is more correlated with transcriptional levels in mammalian genes compared to non-mammalian ones (Fig. 3). For example, in yeast, no clear change is seen for the nucleosomal ΔNPS values in highly (red line) and lowly (blue line) transcribed genes (Fig. 3A-D). By contrast, in mammals, lowly transcribed genes tend to have higher ΔNPS values whereas highly transcribed genes tend to have lower ΔNPS values (Fig. 3E-G). Again, the nucleosome +1 was used for illustration (Supplementary Table S8): the fraction of Type 1 nucleosomes is decreased from lowly transcribed genes (Group 1) to highly transcribed genes (Group 4). By contrast, the fraction of Type 4 nucleosomes is increased from lowly transcribed genes to highly transcribed genes. These results have established a clear link between RNA Pol II transcription levels and the rotational settings of nucleosomes in mammals, that is, nucleosomes are more likely to follow the canonical WW/SS pattern in lowly transcribed genes and the anti-WW/SS pattern in highly transcribed genes.

Overall, we have seen a drastic difference in the distribution of anti-WW/SS nucleosomes in non-mammalian *versus* mammalian genes. That is, anti-WW/SS nucleosomes are enriched in mammalian but not non-mammalian genes. Moreover, the enrichment of anti-WW/SS nucleosomes seems to be associated with RNA Pol II transcription.

### Periodic DNA patterns are missing in mammalian genes

To check whether the observed enrichment of anti-WW/SS nucleosome is caused by underlying DNA patterns, we calculated distance auto-correlation (DAC) functions (Zhurkin 1981; Cui and Zhurkin 2012) of WW dinucleotides in the TSS-surrounding regions (i.e., between −500 bp and +1000 bp relative to TSS). Inspection of the DAC functions in non-mammals reveals a very regular structure, with peaks separated by ∼10 bp (Fig. 4A-C). The same ∼10-bp periodicity is observed in the DAC functions of SS dinucleotides (Supplementary Fig. S11A-C) and the distance cross-correlation (DCC) functions between WW and SS dinucleotides (Supplementary Fig. S12A-C). Note that the DAC functions have the peaks at the distance of ∼10 × n bp (where n = 1, 2, …), whereas the DCC functions have the peaks at the distance of ∼10 × n + 5 bp (where n = 1, 2, …), indicating that WW (or SS) are in phase with each other, whereas WW and SS dinucleotides are out of phase. Our results suggest stable nucleosomes are likely to form with an optimal rotational setting in non-mammalian genes, which may explain small variations of the ΔNPS values observed for nucleosome −1 to +5 (see black lines in Fig. 3A-D).

**Figure 4.**
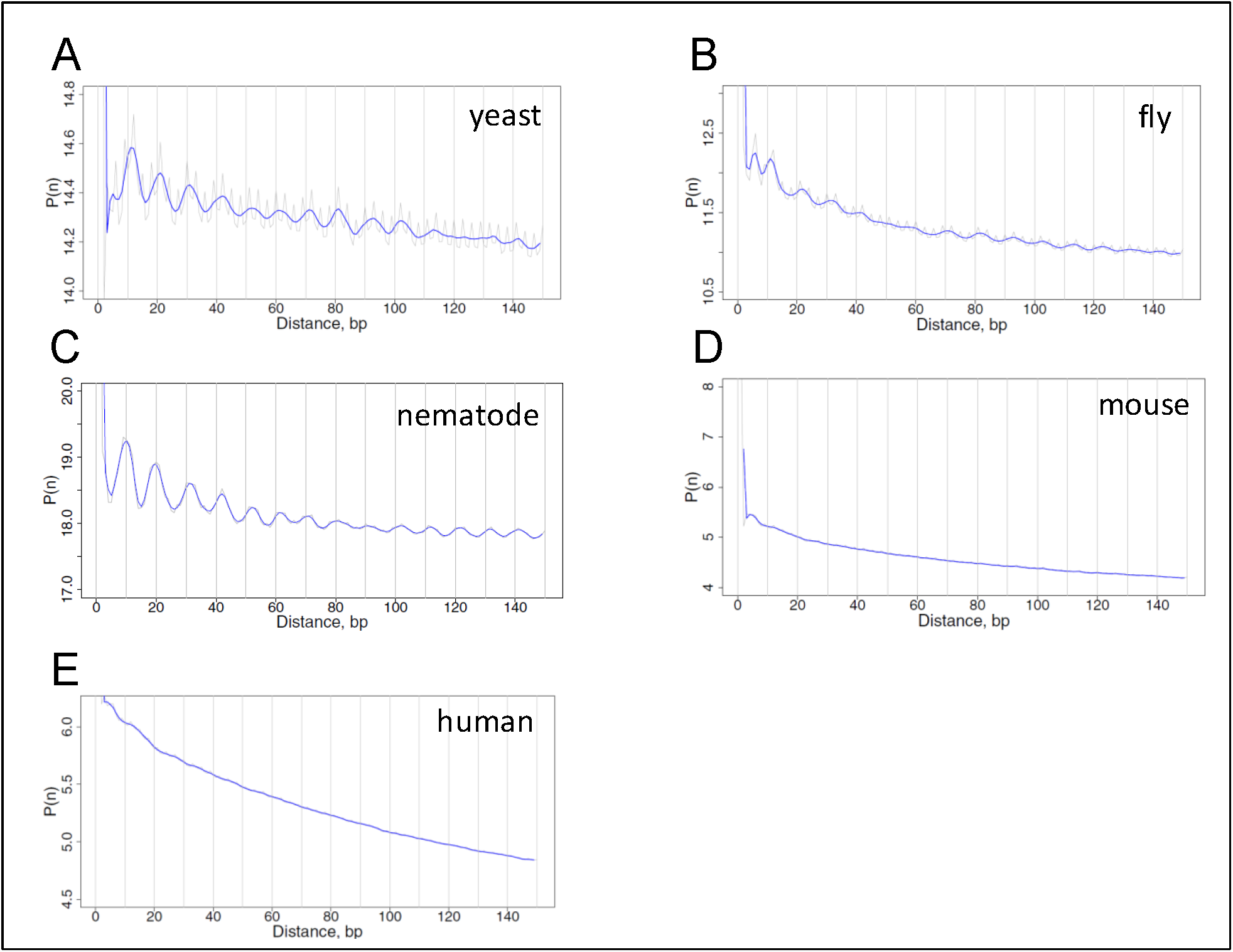
Distance auto-correlation function profiles for WW dinucleotides in yeast (A), fly (B), nematode (C), mouse (D) and human (E) DNA. Genomic fragments [-500 bp, +1000 bp] relative to verified TSSs (position 0) were used for analysis. Both raw (in gray) and 3-bp running average (in blue) values were plotted.

By contrast, both DAC and DCC functions in mouse and human genes exhibit no recognizable peaks (Fig. 4D-E and Supplementary Fig. S11D-E and Fig. S12D-E), indicating that the ∼10-bp periodic WW and SS patterns are missing in mammals, consistent with earlier studies (Tolstorukov et al. 2009; Reynolds et al. 2009). This lack of sequence periodicity in mammals is remarkable because it suggests that the enrichment of anti-WW/SS nucleosomes is not due to intrinsic DNA sequence patterns but to *trans*-acting factors such as RNA Pol II. RNA transcription may facilitate the formation of anti-WW/SS nucleosomes that are intrinsically unstable, which in turn promotes gene transcription (see Discussion).

Taken together, we have observed a significant difference in DNA sequence patterns between mammalian and non-mammalian genes. The non-mammalian DNA exhibits an in-phase pattern of WW (or SS) dinucleotides and an out-of-phase pattern between WW and SS dinucleotides. By contrast, no periodic pattern is observed in mammals. Our data suggest that DNA sequence plays a distinctive role in the rotational settings of nucleosomes in non-mammals *versus* mammals around TSS.

### Anti-WW/SS nucleosomes are not over-represented in mammalian repeats

To examine if the enrichment of anti-WW/SS nucleosomes in mammals is context dependent, we focused on repetitive DNA elements. It is because that up to 40-50% of the murine or human genome contain repeat sequences derived from transposable elements (Waterson et al. 2002), including elements with long terminal repeats (LTR), short interspersed nuclear elements (SINE) and long interspersed nuclear elements (LINE). Many repetitive sequences occur in heterochromatic regions (Grewal and Jia 2007) that are characterized with high levels of condensation throughout the cell cycle (Francastel et al. 2000), low rates of meiotic recombination (Szauter 1984) and the ability to silence gene expression (Eissenberg and Elgin 2000). We therefore hypothesize that the fraction of anti-WW/SS nucleosomes in repeats would differ from that in genic regions (Supplementary Fig. S8).

We first checked the fraction of repeat families in the human genome (Fig. 5A, Supplementary Tables S9-S10) and compared with the fractions of nucleosomes residing in these families (Fig. 5B). Although 18% and 6% of human repeats are LTR and simple repeat elements, only 8% and 1% of NCP fragments are found in these elements respectively, indicating that nucleosomes are depleted in these families, consistent with previous studies (Samans et al. 2014). By contrast, no apparent depletion of nucleosomes is observed in LINE and SINE elements (41% versus 40% for LINE and 29% versus 24% for SINE) (Fig. 5A-B).

**Figure 5.**
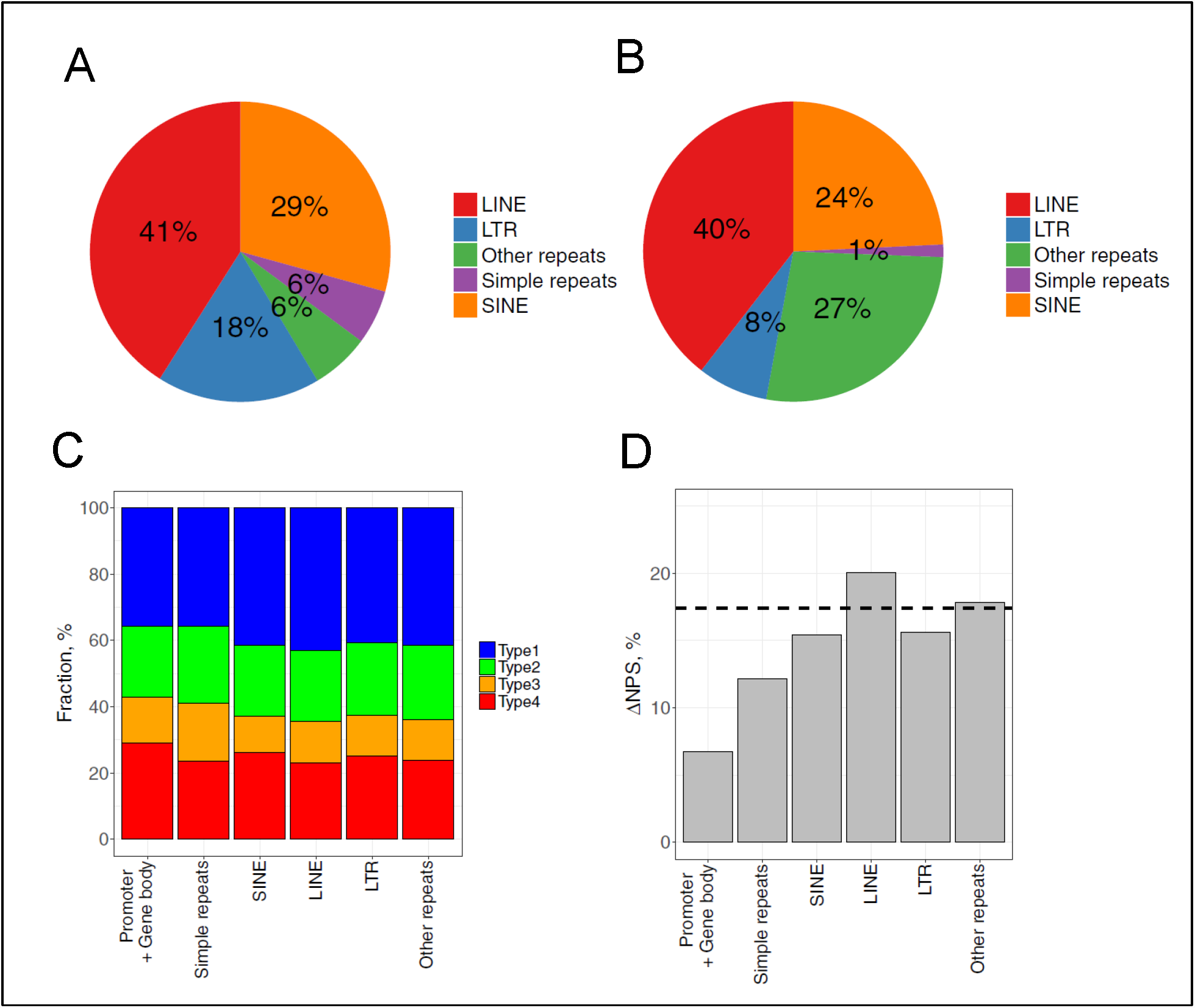
Fractions of DNA sequence patterns and ΔNPS values in human repeat families. (A) Fraction of repeat families in human genome. The fractions were taken from literature (Treangen and Salzberg 2012). (B) Fraction of 147-bp human nucleosomes residing in each repeat family. (C) Fraction of 4 types of nucleosomes in each repeat family. (D) Nucleosome ΔNPS values in genic and repetitive DNA regions. The genomic ΔNPS value is indicated by dashed lines.

Then we calculated the fractions of 4 sequence types and the ΔNPS value across repeat families (Fig. 5C-D) and found that ΔNPS values in all families are higher than that in genic region. In particular, SINE- and LINE-bearing nucleosomes have ΔNPS values close to the genomic value (Fig. 5D). Similar observations are seen in mouse repeats (Supplementary Fig. S13), indicating that anti-WW/SS nucleosomes are not over-represented in mammalian repeats. Examination of the DAC functions of WW dinucleotides in human LINE and SINE elements reveals a ∼10-bp periodicity (Supplementary Fig. S14), which is absent in genic regions (Fig. 4D-E), suggesting that repetitive DNA is organized differently from DNA in genic regions. Because repeat elements often reside in silent chromatin with little or no transcription, it is conceivable that canonical WW/SS nucleosomes are more likely to form in these regions, which may explain why their ΔNPS values are close to the genomic value (Fig. 5D and Supplementary Fig. S13D).

### Chromatin remodelers influence the fraction of anti-WW/SS nucleosomes

Finally, we investigated if other *trans*-acting factors such as chromatin remodeling complexes can influence the abundance of anti-WW/SS nucleosomes. To this end, yeast mutant strains in which single or multiple chromatin remodeling complexes have been deleted were used for analysis. These mutant strains include four single mutants (*isw1Δ, isw2Δ, chd1Δ*, and *rsc8Δ*), three double mutants (*isw1Δ isw2Δ, isw1Δ chd1Δ*, and *isw2Δ chd1Δ*), and one triple mutant (*isw1Δ isw2Δ chd1Δ*) (Ganguli et al. 2014; Ocampo et al. 2016). From the fraction of 4 sequence types (Supplementary Fig. S15) and genomic ΔNPS value (Supplementary Table S12) of each mutant, it is clear that all mutants except *isw2Δ* have significantly different ΔNPS values from the wildtype strain (Student’s t-test *p* < 10^-5^) (Fig. 6), with the average deviation being 1.85% (Supplementary Table S12).

**Figure 6.**
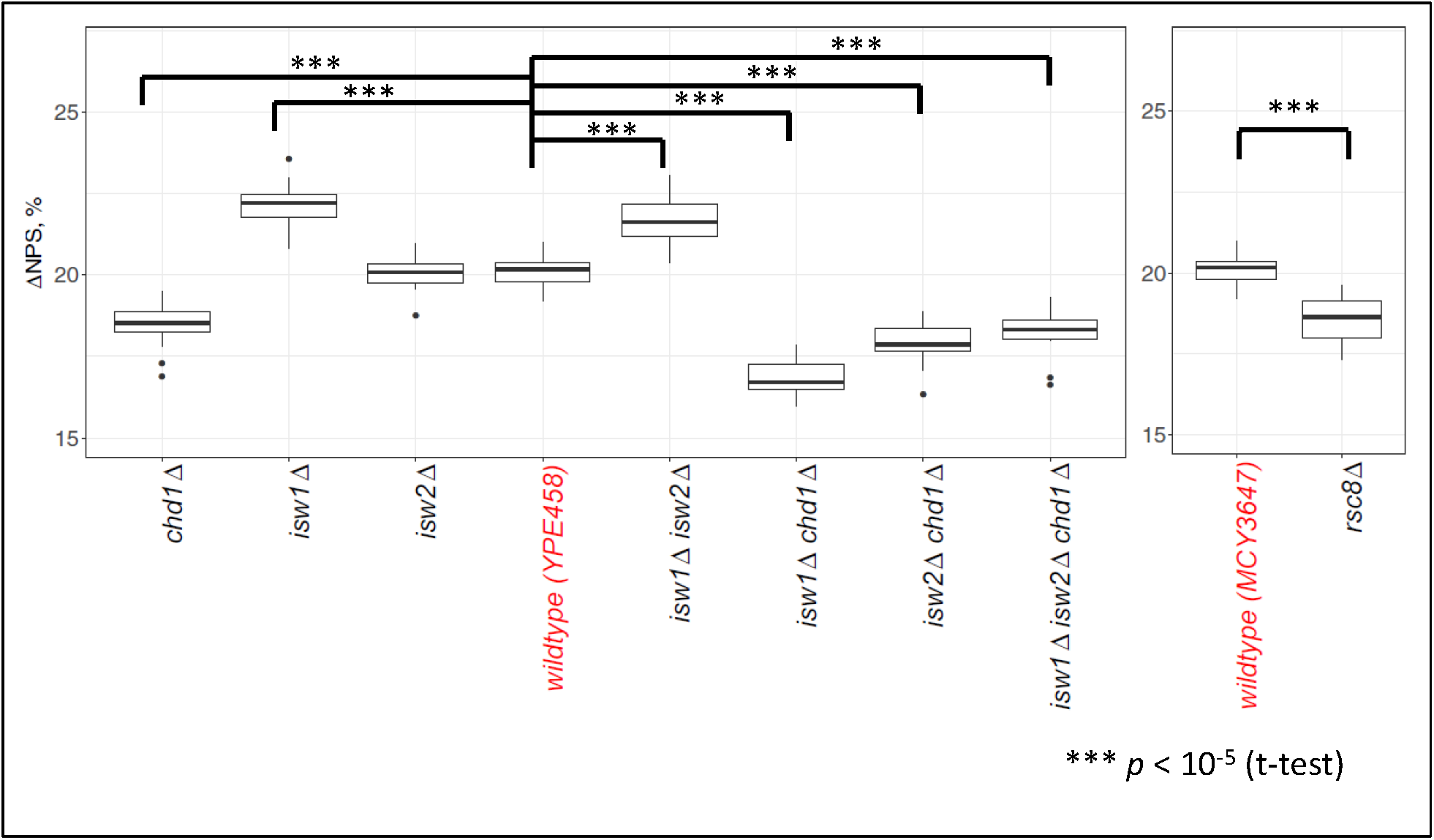
Genomic ΔNPS values in yeast wildtype and mutant strains. ΔNPS values were calculated by chromosomes and shown in a box-whisker plot, with the mean representing the genomic ΔNPS. The genomic ΔNPS values of wildtype strains are used as reference points to compare with those of mutant strains (Supplementary Table S12) in which one or more chromatin remodeler genes are knocked out (Ocampo et al., 2016).

The double mutant strain *isw1Δ chd1Δ* was used for detailed analysis because nucleosome phasing is disrupted (Supplementary Fig. S16A), as shown before (Ocampo et al. 2016). The genomic ΔNPS value of this strain is 16.84%, which is 3.3% smaller than that of the wildtype strain, 20.14% (Supplementary Table S12), suggesting that the deletion of both ISW1 and CHD1 leads to an increase of anti-WW/SS nucleosomes. However, the nucleosome ΔNPS profile around TSS is very similar to that of wildtype strain (Supplementary Fig. S16B and Figs. 3A), indicating that anti-WW/SS nucleosomes are not enriched in the genic regions, although the overall fraction of anti-WW/SS nucleosomes is increased. Similar patterns are seen in other mutants, e.g., *chd1Δ isw1Δ chd1Δ, isw2Δ chd1Δ*, and *isw1Δ isw2Δ chd1Δ* (Fig. 7). Our data suggest that chromatin remodeler(s) have an impact on the fraction but not on the distribution of anti-WW/SS nucleosomes in non-mammals. However, it remains to be determined if this observation holds for mammals.

**Figure 7.**
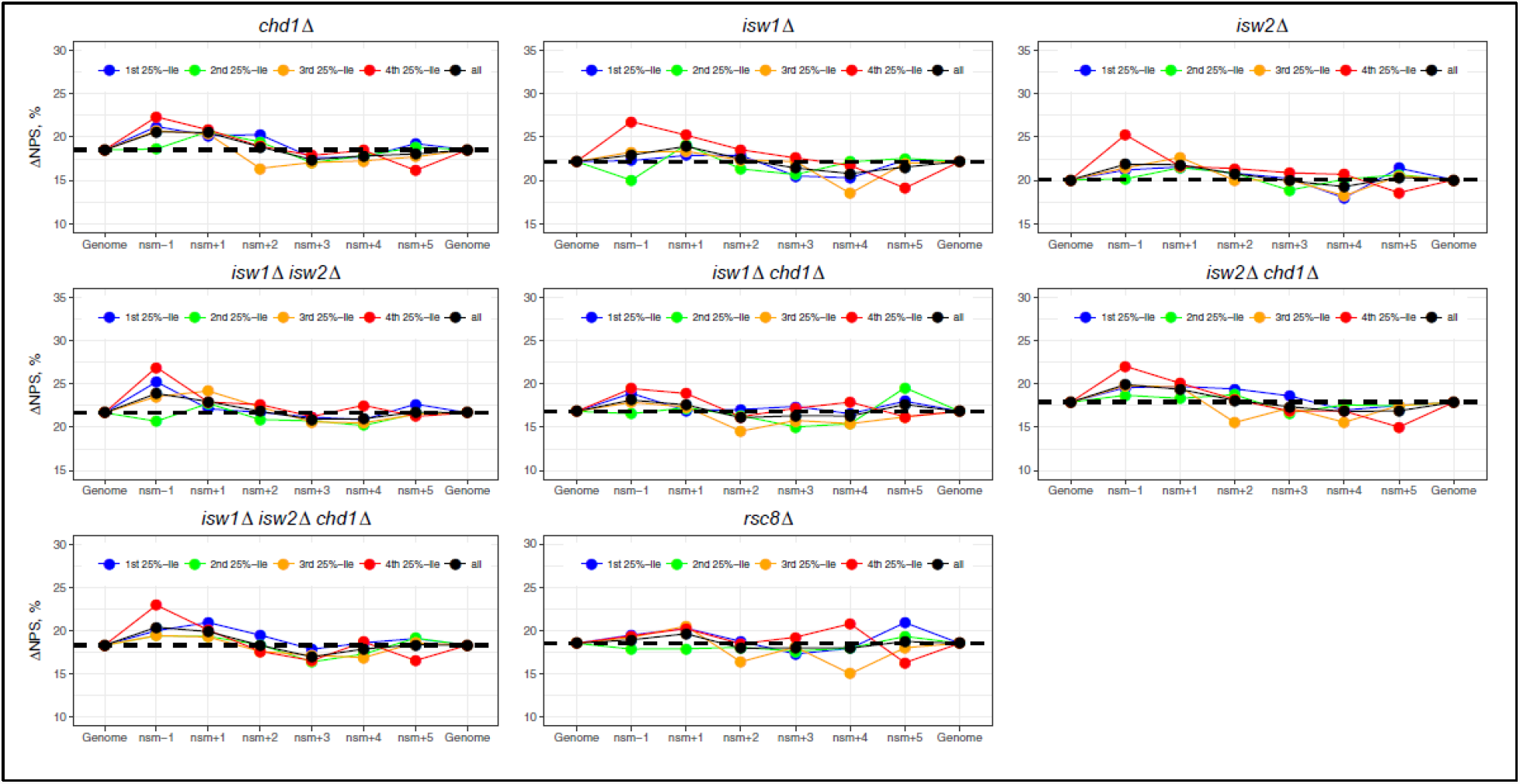
ΔNPS values of promoter and downstream nucleosomes in yeast mutant strains. For the sake of comparison, the ranges of nucleosomes are the same for both the wildtype and mutant strains (Supplementary Table S6). Other notations follow Figure 3.

## Discussions

### A model for *cis* and *trans* determinants of rotational positioning of nucleosomes

We have conducted a comprehensive analysis of WW/SS-based sequence patterns for nucleosomes in five eukaryotes. Our work demonstrates that nucleosomes with the anti-WW/SS pattern are widespread in eukaryotic genomes. Interestingly, these nucleosomes are distributed differently in non-mammals (e.g., yeast, nematodes and fruit flies) *versus* in mammals (e.g., mice and humans). Specifically, anti-WW/SS nucleosomes are enriched in the promoter and genic regions of mammalian genomes but not of non-mammalian genomes (Fig. 8A). We have further analyzed the impact of various *cis*- and *trans*-acting factors on the fraction and the distribution of anti-WW/SS nucleosomes. In light of these findings, we propose a model for the distinct role of *cis* and *trans* factors in the rotational positioning of nucleosomes between non-mammals and mammals (Fig. 8B).

**Figure 8.**
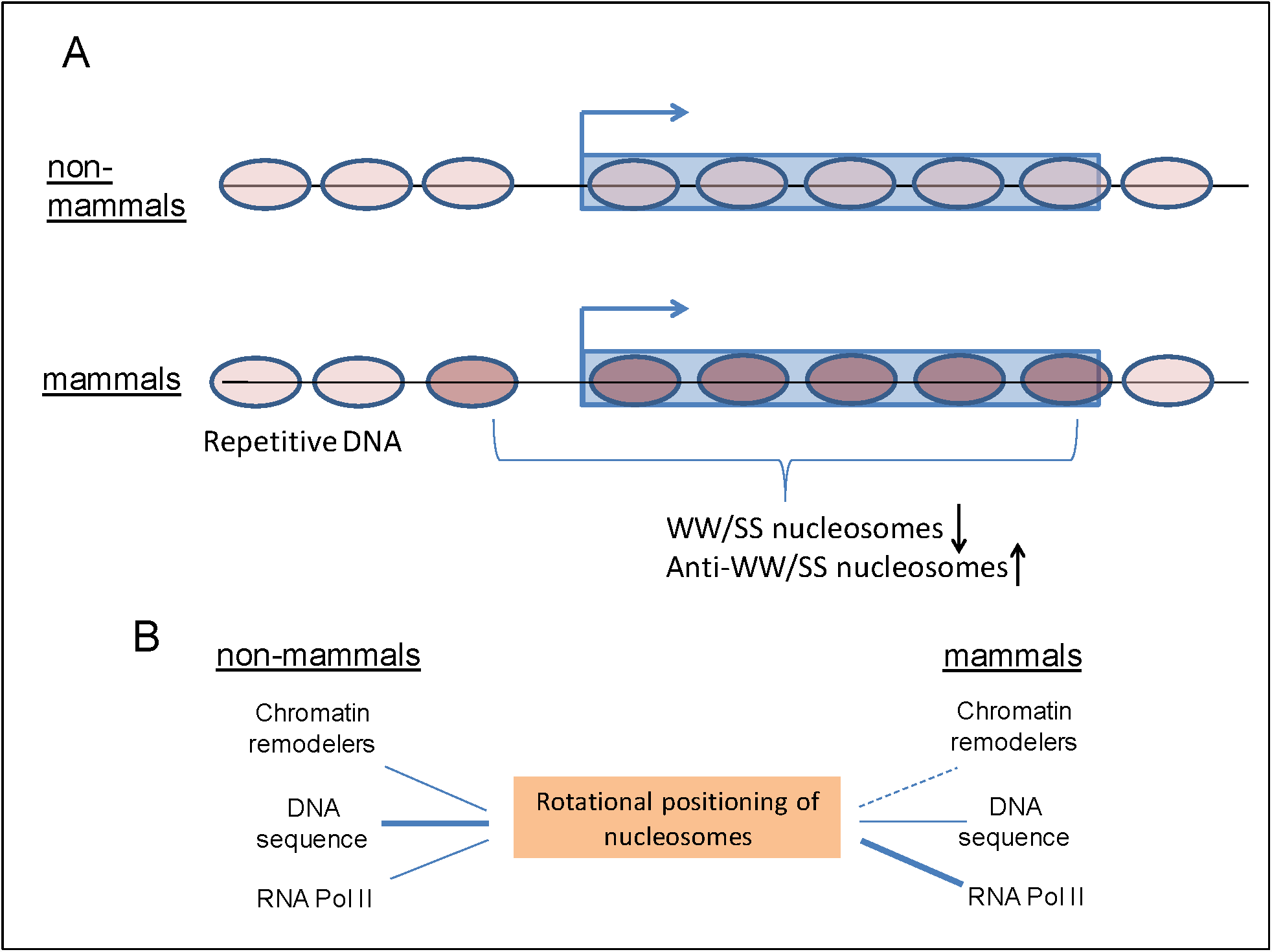
Models for the roles of *cis* and *trans* factors in rotational positioning of nucleosomes. (A) Species-specific distribution of anti-WW/SS nucleosomes. In mammalian genes, the fraction of anti-WW/SS nucleosomes is increased and the fraction of WW/SS nucleosomes is decreased. This change is not seen in non-mammalian genes. Nucleosomes −1 to +5 in mammals are depicted in dark brown and other nucleosomes (including those residing in mammalian repetitive DNA) are depicted in light brown. (B) Distinct role of *cis* and *trans* factors in rotational positioning of nucleosomes. Both cis-acting factors (e.g., DNA sequences) and trans-acting factors (e.g., chromatin remodeling complexes and RNA Pol II) can affect rotational settings of nucleosomes. In non-mammals, DNA sequence play a more important role than RNA Pol II in rotational positioning because (1) the ∼10-bp periodic WW (or SS) patterns are pronounced (represented by thick lines); (2) no clear change in the ΔNPS values between highly and lowly transcribed genes. By contrast, in mammalian genes, RNA Pol II plays a more important role (represented by a thick lines) because (1) the ∼10-bp periodicity is diminished, and (2) a clear change in ΔNPS values occurs between highly and lowly transcribed genes. Chromatin remodelers have an impact on the rotational setting of nucleosomes in non-mammals, but this effect is unclear in mammals (represented by dashed lines).

Based on our model, the rotational positioning of nucleosomes can be influenced by *cis*- and *trans*-acting factors such as DNA sequence, chromatin remodeling complexes, and RNA Pol II in both non-mammals and mammals. However, the relative importance of these factors differ. In non-mammals, the *cis*-acting factors play a more important role than *trans*-acting factors because (1) genomic DNA exhibits sequence patterns with a ∼10-bp periodicity (Fig. 4A-C and Supplementary Figs. S11A-C and S12A-C), and (2) the fraction of anti-WW/SS nucleosomes varies little across the genomes (Fig. 3A-D and Fig. 7). By contrast, the *trans*-acting factors such as RNA Pol II appears to be more important in mammals, due to a lack of periodic sequence patterns (Fig. 4D-E and Supplementary Figs. S11D-E and S12D-E), and the enrichment of anti-WW/SS nucleosomes in mammalian promoters and genic regions (Fig. 3E-G).

It is worth noting that overall, periodic WW (or SS) patterns appear to be very strong in yeast, become weaker in nematodes and flies, and completely disappear in mice and humans (Fig. 4 and Supplementary Figs. S11 and S12). This implies that the importance of *cis*-acting factors in the rotational positioning of nucleosomes decreases in evolution, especially in promoter and genic regions. However, the periodic sequence patterns are preserved in mammalian repetitive DNA elements (Supplementary Fig. S14) and anti-WW/SS nucleosomes are not over-represented (Fig. 5D). These data suggest that molecular mechanisms governing the rotational settings of nucleosomes in mammals are context dependent.

### Implications of anti-WW/SS nucleosomes for RNA Pol II transcription

The interactions between RNA Pol II transcription and nucleosomes have been extensively studied since the discovery of nucleosomal barrier (Studitsky et al. 1995; Teves et al. 2014). Early work has established that the nucleosome presents a strong barrier to transcription *in vitro*, and the nature of this barrier has been studies by structural, biochemical and biophysical studies (Gaykalova et al. 2015; Hall et al 2009). However, the questions of how the Pol II machinery efficiently overcomes nucleosomes and transcribes through gene body *in vivo* remain unclear. Various *trans* factors modulating the nucleosomal barrier have been proposed (Teves et al. 2014), including histone modifications, histone variant replacement, histone chaperones and nucleosome remodelers. Our work suggests that *cis* factors such as DNA sequence also contribute to modulation of nucleosomal barrier, especially in mammals.

As shown above, anti-WW/SS nucleosomes are enriched in promoters and genic regions of mammals, and this enrichment is increased with the transcriptional levels. One possible explanation is that because of the intrinsic instability of nucleosomes with respect to rotational settings, RNA Pol II transcription is promoted. However, the formation of anti-WW/SS nucleosomes is not energetically favorable because GC-containing dinucleotides in the minor-GBS do not have favorable interactions with the “sprocket” arginine residues in histones (see Introduction). In addition, the enrichment of this type of nucleosomes is not seen in highly transcribed genes in non-mammals (Fig. 3A-D). An alternative explanation is that RNA Pol II transcription helps to position anti-WW/SS nucleosomes that are thermodynamically unstable. In these nucleosomes, the presence of SS dinucleotides in the minor-GBS weakens DNA-histone interactions, which may facilitate the unwrapping of DNA from nucleosomes (Ngo et al. 2016), thereby promoting RNA Pol II transcription. This tendency may also help explain histone eviction observed in highly transcribed genes (Kristjuhan and Svejstrup 2004; Schwabish and Struhl 2004).

### Rotational positioning and binding of transcriptional factors in mammalian repeats

Recent studies have shown that nucleosomes are depleted in certain types of repeat elements (e.g., simple repeats and LTR), but not in other types (e.g., SINE and LINE) in human sperms (Samans et al. 2014). Indeed, LTR/ERV elements are highly enriched in open chromatin regions (Jacques et al. 2013), whereas Alu elements that belong to the SINE superfamily harbor two nucleosomes (Englander et al. 1993; Englander and Howard 1995; Tanaka et al. 2010). Our analysis of the human nucleosome data, which were derived from lymphoblastoid cell lines (Gaffney et al. 2012), is in general agreement with these observations in sperms (Fig. 5A-B). A group of TFs known as pioneer factors (e.g., p53, Oct4, Sox2, Klf4 etc.) is able to interact with nucleosomal DNA (Saho et al. 2009; Laptenko et al. 2011; Soufi et al. 2015) and many of them preferentially bind to repetitive DNA elements (Kunarso et al. 2010).

We and other groups have shown that the rotational positioning of nucleosomes is critical for TF binding including p53 (Sahu et al. 2009; Cui and Zhurkin 2014). We have found that an Alu-residing nucleosome, if taking the optimal rotational setting, help to expose the putative binding sites of p53 (Cui and Zhurkin 2011). The present study has demonstrated that nucleosomes are likely to take the optimal rotational setting in human repeats including SINE/Alu elements (Fig. 8D), suggesting that many p53 binding sites in Alu elements are likely to be exposed. This is also in agreement with our recent findings that a large amount (∼40%) of p53 binding sites occurs in various repeats including Alu (Bao et al. 2017). Further analyses are needed to address the biological roles of anti-WWSS nucleosomes in repeats to see if, in these cases, TF binding sites are buried inside of nucleosomes and inhibitory for TF binding, as we suggested for p53 (Cui and Zhurkin 2014). Understanding the rotational positioning of nucleosomes in human repeats will shed new lights on mechanisms regulating retrotransposon activities in a normal or diseased cellular context.

## Materials and Methods

### Definition of Minor- and major-groove bending sites in a nucleosomal DNA fragment

Minor- and major-groove bending sites (GBS) in a 147-bp nucleosomal DNA fragment were defined previously (Cui and Zhurkin 2010). Briefly, minor-GBS are located at the superhelical locations (SHL) ±0.5, ±1.5, …, ±6.5, while major-GBS are located at SHL ±1, ±2, …, ±6. Each site is 3 or 4 base pairs in length. In total, there are 14 minor-GBS and 12 major-GBS along a 147-bp NCP fragment. Note that the minor-GBS at SHL ±0.5 (shown in grey in Supplemental Figure S1) are not included for analysis because out-of-phase WW/SS peaks were observed at these two sites (Satchwell et al. 1986). As a result, 12 minor-GBS covering 48 bp, and 12 major-GBS covering 44 bp are used in our analysis (Supplemental Table S1). Since each minor-GBS (4 bp in length) holds 3 unique dinucleotides, the maximum number of dinucleotides in 12 minor-GBS is 36 (=3 × 6 × 2). Similarly, the maximum number of dinucleotides in 12 major-GBS is 32 (= (3 × 4 + 2 × 2) × 2). To account for this difference, we assign a coefficient of 1.125 (=36/32) to dinucleotides occurring at major-GBS (Table 1).

### Classification of nucleosomes with different NPS patterns

Nucleosomal DNA sequences are divided into 4 types based on relative abundance of WW and SS dinucleotides in minor- and major-GBS (Table 1). Type 1 nucleosomes have the canonical WW/SS pattern (i.e., WW are more abundant in minor-GBS than in major-GBS, whereas SS are more abundant in major-GBS than in minor-GBS). Specifically, for a given 147-bp NCP fragment, if the total number of WW dinucleotides in the 12 minor-GBS is greater than its counterpart in the 12 major-GBS, and the total number of SS dinucleotides in the 12 major-GBS is greater than its counterpart in the 12 minor-GBS, this fragment is classified as Type 1 nucleosomal DNA (Fig. 1A).

Type 4 nucleosomes are characterized with an anti-WW/SS pattern. That is, if a 147-bp NCP fragment has more WW dinucleotides in 12 major-GBS than in minor-GBS, and has more SS dinucleotides in minor-GBS than in major-GBS, this fragment is classified as Type 4 nucleosomal DNA (Fig. 1D). Type 2 and 3 nucleosomes have ‘mixed’ patterns, in which both WW and SS are abundant in minor-GBS (Type 2, Fig. 1B) or in major-GBS (Type 3, Fig. 1C). Specifically, a Type-2 NCP fragment has more WW and SS dinucleotides in minor-GBS than in major-GBS, while a Type-3 NCP fragment has more WW and SS dinucleotides in major-GBS than in minor-GBS. For a given genomic region, the difference between Type 1 and Type 4 nucleosomes in percentage (%) is denoted as ΔNPS. That is, ΔNPS = Type 1 (%) – Type 4 (%).

### High-resolution nucleosome datasets in yeast and higher eukaryotes

A total of 18 high-resolution nucleosomal DNA datasets from various eukaryotes were used in this study (Supplemental Table S2), in which nucleosome positions were mapped by MNase-Seq or the chemical methods. Two datasets, one from yeast (Brogaard et al. 2012) and the other from mouse (Voong et al. 2016), were generated by the chemical method. Sixteen datasets were produced by paired-end MNase-Seq. Because this sequencing technique generates short reads on both ends of a nucleosomal DNA fragment, after mapping them to the reference genome, the length of the fragment can be precisely determined. Only the fragments of 147 bp in length were analyzed in this study (Supplemental Table S3). This is because that in these cases, minor- and major-GBS can be determined in an unambiguous manner based on a high-resolution nucleosome crystal structure (Cui and Zhurkin 2010, also see Supplemental Fig. 1).

Paired-end nucleosome sequence data from yeast include wildtype (WT) cells YPE458 (Ocampo et al. 2016), and MCY3647 (Ganguli et al. 2014) and mutant (MT) cells (Ganguli et al. 2014; Ocampo et al. 2016) including single mutants (*isw1Δ, isw2Δ, chd1Δ, rsc8Δ*), double mutants (*isw1Δ isw2Δ, isw1Δ chd1Δ, isw2Δ chd1Δ*), and triple mutants (*isw1Δ isw2Δ chd1Δ*). The paired reads were aligned to the *Saccharomyces cerevisiae* genome using ELAND (Illumina). Only the reads uniquely aligned to the genome with no mismatch were selected. Note that all yeast strains (wildtype or mutants) were grown to mid-log phase in synthetic complete (SC) medium. All of the wildtype strains (YDC111, YPE458 and MCY3647) should be equivalent for most purposes, since they grew in the same SC medium.

Paired-end MNase-seq data were also taken from *C. elegans* embryos and sperms (Tabuchi et al. 2018), *Drosophila* S2 cells (Chereji et al. 2016, Fuda and Lis, 2015), mouse embryonic stem cells (mESC) (Voong et al. 2016), and human lymphoblastoid cell lines 18508 and 19238 (Gaffney et al. 2012). BAM files were either downloaded from NCBI GEO database or obtained by mapping raw reads to the corresponding genomes using the default setting of Bowtie (Langmead et al. 2009).

### Genome-wide gene expression (RNA-seq) data in yeast and humans

The RPKM (<u>R</u>eads <u>P</u>er <u>K</u>ilobase of transcript per <u>M</u>illion) values in RNA-seq data of *C. elegans* embryos (GSM1652723) and sperms (GSM3188165), Drosophila S2 cells (GSM410195), mouse mESC (GSM2183915) and human granulocytes (GSM678066) were used to divide genes into 4 groups based on transcriptional frequencies (Supplementary Table S5).

The RNA-seq data for the SC condition were used for analysis (Waern and Snyder 2013). Raw reads were mapped to the yeast genome by the default setting of Bowtie (Langmead et al. 2009). The RPKM values of the yeast data were calculated following the same method as the human data (Valouev et al. 2010).

### Repeat elements in the human genome

Mouse (mm10) and human (hg19) repetitive region positions were downloaded from the UCSC Genome Browser. The repeat elements were identified using RepeatMasker (v3.2.7) and Repbase Update (9.11). The major types of repeat elements were selected for analysis, including SINE (Alu, MIR), LINE (CR1, L1, L2 and RTE), LTR (ERV1, ERVK, ERVL and Gypsy), Simple Repeat ((TG)_n_, (TCG)_n_, (CACTC)_n_, (GAGTG)_n_, and (TATATG)_n_). The remaining repeat elements were included into the “Other” category including MuDR, PiggyBac, TcMar-Mariner, hAT-Charlie. Only the elements with >150 bp in length (i.e., approximately the size of one nucleosome) were selected for analysis.

Since human nucleosomes were mapped to the genome assembly hg18 (Gaffney et al. 2012), the UCSC LiftOver utility was used along with the hg18-hg19 chain file to convert the hg19 repeat elements to their corresponding positions in human genome assembly hg18 (Supplemental Table S10). Nucleosomes overlapped with these repeat families were summarized in Supplemental Table S11.

### Distance auto- and cross-correlation function

The distance auto-correlation (DAC) and distance cross-correlation (DCC) functions have been discussed previously (Zhurkin 1981; Cui et al. 2012). In the present study, DAC function was used to calculate the correlation between WW or SS dinucleotides. That is, if a WW dinucleotide (AA, TT, AT or TA) is separated from another WW dinucleotide by a distance *d*, one occurrence is counted for that distance in the DAC function. For the DCC function, we counted how many times a WW dinucleotide is separated from a SS dinucleotide (GG, CC, GC or CG) by a distance *d*, where *d* is between 1 and 150 bp.

## References

Baldi P, Brunak S, Chauvin Y, Krogh A. 1996. Naturally occurring nucleosome positioning signals in human exons and introns. J Mol Biol 263: 503–510.

Bao F, LoVerso PR, Fisk JN, Zhurkin VB, Cui F. 2017. P53 binding sites in normal and cancer cells are characterized by distinct chromatin context. Cell Cycle 16: 2073–2085.

Brogaard K, Xi L, Wang JP, Widom J. 2012. A map of nucleosome positions in yeast at base-pair resolution. Nature 486: 496–501.

Calladine CR, Drew HR. 1986. Principles of sequence-dependent flexure of DNA. J Mol Biol 192: 907–918.

Cui F, Zhurkin VB. 2009. Distinctive sequence patterns in metazoan and yeast nucleosomes: implications for linker histone binding to AT-rich and methylated DNA. Nucleic Acids Res. 37: 2818–2829.

Cui F, Zhurkin VB. 2010. Structure-based analysis of DNA sequence patterns guiding nucleosome positioning in vitro. J Biomol Struct Dyn 27: 821–841.

Cui F, Cole HA, Clark DJ, Zhurkin VB. 2012. Transcriptional activation of yeast genes disrupt intragenic nucleosome phasing. Nucleic Acids Res 40: 10753–10764.

Cui F, Sirotin MV, Zhurkin VB. 2011. Impact of Alu repeats on the evolution of human p53 binding sites. Biol Direct 6:2

Cui F, Chen L, LoVerso PR, Zhurkin VB. 2014. Prediction of nucleosome rotational positioning in yeast and human genomes based on sequence-dependent DNA anisotropy. BMC Bioinformatics 15: 313.

Cui F, Zhurkin VB. 2014. Rotational positioning of nucleosomes facilitates selective binding of p53 to response elements associated with cell cycle arrest. Nucleic Acids Res 42: 836–847.

Chereji RV, Kan TW, Grudniewska MK, Romashchenko AV, Berezikov E, Zhimulev IF, Guryev V, Morozov AV, Moshkin YM. 2016. Genome-wide profiling of nucleosome sensitivity and chromatin accessibility in Drosophila melanogaster. Nucleic Acids Res 44: 1036–1051.

Chereji RV, Ramachandran S, Bryson TD, Henikoff S. 2018. Precise genome-wide mapping of single nucleosomes and linkers in vivo. Genome Biol. 19: 19.

Davey CA, Sargent DF, Luger K, Maeder AW, Richmond TJ. 2002. Solvent mediated interactions in the structure of the nucleosome core particle at 1.9Å resolution. J Mol Biol 391: 1097–1113.

Dingwall C, Lomonossoff GP, Laskey RA. 1981. High sequence specificity of micrococcal nuclease. Nucleic Acids Res. 9: 2659–2673.

Drew HR, Travers AA. 1985. DNA bending and its relation to nucleosome positioning. J Mol Biol 186: 773–790.

Eissenberg JC, Elgin SC. 2000. The HP1 protein family: getting a grip on chromatin. Curr Opin Genet Dev 10: 204–210.

Englander EW, Howard BH 1995. Nucleosome positioning by human Alu elements in chromatin. J Biol Chem 270: 10091–10096.

Englander EW, Wolffe AP, Howard BH. 1993. Nucleosome interactions with a human Alu element. J Biol Chem 268: 19565–19573.

Field Y, Kaplan N, Fondufe-Mittendorf Y, Moore I, Sharon E, Lubling Y, Widom J, Segal E. 2008. Distinct modes of regulation by chromatin encoded through nucleosome positioning signals. PLoS Comput. Biol. 4: e1000175

Flaus A, Luger K, Tan S, Richmond TJ. 1996. Mapping nucleosome position at single base-pair resolution by using site-directed hydroxyl radicals. Proc. Natl. Acad. Sci. USA. 93: 1370–1375

Flaus A, Richmond TJ. 1998. Positioning and stability of nucleosomes on MMTV 3’LTR sequences. J Mol. Biol. 275: 427–441.

Francastel C, Schubeler D, Martin DI, Groudine M. 2000. Nuclear compartmentalization and gene activity. Nat. Rev. Mol. Cell Biol. 1: 137–143.

Gaffney DJ, McVicker G, Pai AA, Fondufe-Mittendorf YN, Lewellen N, Michelini K, Widom J, Gilad Y, Pritchard JK. 2012. Controls of nucleosome positioning in the human genome. PLoS Genet 8: e1003036.

Ganguli D, Chereji RV, Iben JR, Cole HA, Clark DJ. 2014. RSC-dependent constructive and destructive interference between opposing arrays of phased nucleosomes in yeast. Genome Res 24: 1637–1649.

Gaykalova DA, Kulaeva OI, Volokh O, Shaytan AK, Hsieh F-K, Kirpichnikov MP, Sokolova OS, Studitsky VM. 2015. Structural analysis of nucleosomal barrier to transcription. Proc. Natl. Acad. Sci. USA. 112: E5787–E5795.

Grewal SIS and Jia S. 2007. Heterochromatin revisited. Nature Rev Genet 8: 35–46.

Hall MA, Shundrovsky A, Bai L, Fulbright RM, Lis JT, Wang MD. 2009. High-resolution dynamic mapping of histone-DNA interactions in a nucleosome. Nat Struct Mol Biol. 16: 124–129.

Horz W, Altenburger W. 1981. Sequence specific cleavage of DNA by micrococcal nuclease. Nucleic Acids Res. 9: 2643–2658.

Ioshikhes I, Bolshoy A, Trifonov EN. 1992. Preferred positions of AA and TT dinucleotides in aligned nucleosome DNA sequences. J Biomol Struct Dyn 9: 1111–1117.

Ioshikhes I, Bolshoy A, Derenshteyn K, Borodovsky M, Trifonov EN. 1996. Nucleosome DNA sequence pattern revealed by multiple alignment of experimentally mapped sequences. J Mol Biol 262: 129–139.

Ioshikhes I, Hosid S, Pugh BF. 2011. Variety of genomic DNA patterns for nucleosome positioning. Genome Res 21: 1863–1871.

Jiang C, Pugh BF. 2009. A compiled and systematic reference map of nucleosome positions across the Saccharomyces cerevisiae genome. Genome Biol 10: R109.

Laptenko O, Beckerman R, Freulich E, Prives C. 2011. p53 binding to nucleosomes within the p21 promoter in vivo leads to nucleosome loss and transcriptional activation. Proc Natl Acad Sci USA 108: 10385–10390.

Jacques PE, Jeyakani J, Bourque G. 2013. The majority of primate-specific regulatory sequences are derived from transposable elements. PLoS Genet. 9: e1003504.

Kassabov SR, Henry NM, Zofall M, Tsukiyama T, Bartholomew B. 2002. High-resolution mapping of changes in histone-DNA contacts of nucleosomes remodeled by ISW2. Mol. Cell Biol. 22: 7524–7534.

Kogan SB, Kato M, Kiyama R, Trifonov EN. 2006. Sequence structure of human nucleosome DNA. J Biomol Struct Dyn 24: 43–48.

Kristjuhan A, Svejstrup JQ. 2004. Evidence for distinct mechanisms facilitating transcription elongation through chromatin in vivo. EMBO J 23: 4243–4252.

Kunarso G, Chia NY, Jeyakani J, Hwang C, Lu X, Chan YS, Ng HH, Bourque G. 2010. Transposable elements have rewired the core regulatory network of human embryonic stem cells. Nat Genet 42: 631–634.

Langmead B, Trapnell C, Pop M, Salzberg SL. 2009. Ultrafast and memory-efficient alignment of short DNA sequences to the human genome. Genome Biol 10: R25.

Lowary PT, Widom J. 1998. New DNA sequence rules for high affinity binding to histone octamer and sequence-directed nucleosome positioning. J Mol Biol 276: 19–42.

Lu Q, Wallrath LL, Elgin SC. 1994. Nucleosome positioning and gene regulation. J Cell Biochem 55: 83–92.

Luger K, Mader AW, Richmond RK, Sargent DF, Rochmond TJ. 1997. Crystal structure of the nucleosome core particle at 2.8Å resolution. Nature 389: 251–260.

Mavrich TN, Ioshikhes IP, Venters BJ, Jiang C, Tomsho LP, Qi J, Schuster SC, Albert I, Pugh BF. 2008. A barrier nucleosome model for statistical positioning of nucleosomes throughout the yeast genome. Genome Res. 18: 1073–1083.

Mengeritsky G, Trifonov EN. 1983. Nucleotide sequence-directed mapping of the nucleosomes. Nucleic Acids Res 11: 3833–3851.

Moyle-Heyrman G, Zaichuk T, Xi L, Zhang Q, Uhlenbeck OC, Holmgren R, Widom J, Wang JP. 2013. Chemical map of Schizosaccharomyces pombe reveals species-specific features in nucleosome positioning. Proc. Natl. Acad. Sci. USA 110: 20158–20163.

Muthurajan UM, Bao Y, Forsberg LJ, Edayathumangalam RK, Suto RK, Chakravarthy S. Dyer PN, Luger K. 2003. Structure and dynamics of nucleosomal DNA. Biopolymers 68: 547–556.

Ngo TTM, Zhang Q, Zhou R, Yodh JG, Ha T. 2016. Asymmetric unwrapping of nucleosomes under tension directed by DNA local flexibility. Cell 160: 1135–1144.

Ocampo J, Chereji RV, Eriksson PR, Clark DJ. 2016. The ISW1 and CHD1 ATP-dependent chromatin remodelers compete to set nucleosome spacing in vivo. Nucleic Acids Res 44: 4625–4635.

Panetta G, Buttinelli M, Flaus A, Richmond TJ, Rhodes D. 1998. Differential nucleosome positioning on Xenopus oocyte and somatic 5S RNA genes determines both TFIIIA and H1 binding: a mechanism for selective H1 repression. J. Mol. Biol. 282: 683–697.

Reynolds SM, Bilmes JA, Noble WS. 2009. On the Relationship between DNA Periodicity and Local Chromatin Structure. In: Batzoglou S. (eds) Research in Computational Molecular Biology.

RECOMB 2009. Lecture Notes in Computer Science, vol. 5541. Springer, Berlin, Heidelberg

Rohs R, West SM, Sosinsky A, Liu P, Mann RS, Honig B. 2009. The role of DNA shape in protein-DNA recognition. Nature 461: 1248–1253.

Sahu G, Wang D, Chen CB, Zhurkin VB, Harrington RE, Appella E, Hager GL, Nagaich AK. 2009. p53 binding to nucleosomal DNA depends on the rotational positioning of DNA response element. J Biol Chem 285:1321–1332.

Samans B, Yang Y, Krebs S, Sarode GV, Blum H, Reichenbach M, Wolf E, Steger K, Sansranjavin T, Schagdarsurengin U. 2014. Uniformity of nucleosome preservation pattern in mammalian sperm and its connection to repetitive DNA elements. Dev Cell 30: 23–35.

Satchwell SC, Drew HR, Travers AA. 1986. Sequence periodicities in chicken nucleosome core DNA. J Mol Biol 191: 659–675.

Schwabish MA, Struhl K. 2004. Evidence for eviction and rapid deposition of histones upon transcriptional elongation by RNA polymerase II. Mol Cell Biol 24: 10111–10117.

Sekinger EA, Moqtaderi Z, Struhl K. 2005. Intrinsic histone-DNA interactions and low nucleosome density are important for preferential accessibility of promoter regions in yeast. Mol Cell 18: 736–748.

Shivaswamy S, Bhinge A, Zhao Y, Jones S, Hirst M, Iyer VR. 2008. Dynamic remodeling of individual nucleosomes across a eukaryotic genome in response to transcriptional perturbation. PLoS Biol 6: e65.

Soufi A, Garcia MF, Jaroszewicz A, Osman N, Pellegrini M, Zaret KS. 2015. Pioneer transcription factors target partial DNA motifs on nucleosomes to initiate reprogramming. Cell 161: 555–568.

Struhl K, Segal E. 2013. Determinants of nucleosome positioning. Nat Struct Mol Biol 20: 267–273.

Sullivan SA and Landsman D. 2003. Characterization of sequence variability in nucleosome core histone folds. Proteins 52: 454–465.

Szauter P. 1984. An analysis of regional constrains on exchange in Drosophila melanogaster using recombination-defective meiotic mutants. Genetics 106: 45–71.

Tabuchi TM, Rechtsteiner A, Jeffers TE, Egelhofer TA, Murphy CT, Strome S. 2018. Caenorhabditis elegans sperm carry a histone-based epigenetic memory of both spermatogenesis and oogenesis. Nat Commun 9: 4310.

Tanaka Y, Yamashita R, Suzuki Y, Nakai K. 2010. Effects of Alu elements on global nucleosome positioning in the human genome. BMC Genomics 11: 309.

Tolstorukov MY, Kharchenko PV, Goldman JA, Kingston RE, Park PJ. 2009. Comparative analysis of H2A.Z nucleosome organization in the human and yeast genomes. Genome Res 19: 967–977.

Treangen TJ, Salzberg SL. 2012. Repetitive DNA and next-generation sequencing: computational challenges and solutions. Nat Rev Genet 13: 36–46.

Trifonov EN. 2011. Thirty years of multiple sequence codes. Genomics Proteomics Bioinformatics 9: 1–6.

Uberbacher EC, Harp JM, Bunick GJ. 1988. DNA sequence patterns in pre*cis*ely positioned nucleosome. J Biomol Struct Dyn 6: 105–120.

Valouev A, Johnson SM, Boyd SD, Smith CL, Fire AZ, Sidow A. 2011. Determinants of nucleosome organization in primary human cells. Nature 474: 516–520.

Voong LN, Xi L, Sebeson AC, Xiong B, Wang JP, Wang X. 2016. Insights into nucleosome organization in mouse embryonic stem cells through chemical mapping. Cell 167: 1555–1570.

Waern K, Snyder M. 2013. Extensive transcript diversity and novel upstream open reading frame regulation in yeast. G3 Genes Genom Genet 3: 343–352.

Wang D, Ulyanov NB, Zhurkin VB. 2010. Sequence-dependent Kink-and-Slide deformations of nucleosomal DNA facilitated by histone arginines bound in the minor groove. J Biomol Struct Dyn 27: 843–859.

Waterson RH et al. 2002. Initial sequencing and comparative analysis of the mouse genome. Nature 420: 520–562.

West SM, Rohs R, Mann RS, Honig B. 2010. Electrostatic interactions between arginines and the minor groove in the nucleosomes. J Biomol Struct Dyn 27: 861–866.

Yuan G, Liu Y, Dion MF, Slack MD, Wu LF, Altschuler SJ, Rando OJ. 2005. Genome-scale identification of nucleosome positions in *S. cerevisiae*. Science 309: 626–630.

Zhurkin VB. 1981. Periodicity in DNA primary structure is defined by secondary structure of the coded protein. Nucleic Acids Res 9: 1963–1971.

Zhurkin VB. 1983. Specific alignment of nucleosomes on DNA correlates with periodic distribution of purine-pyrimidine and pyrimidine-purine dimers.

