## Supplementary Figures for "An unusual nucleosomal sequence pattern is enriched in mammalian genes"

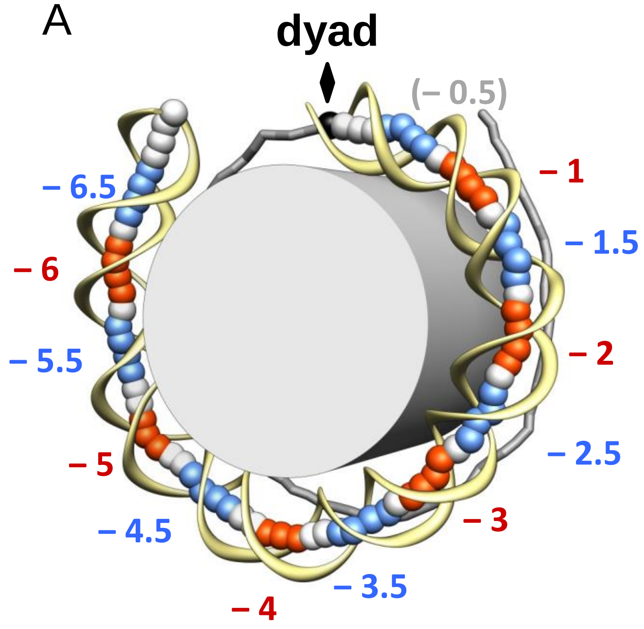

B

**minor-groove bending sites**

| DNA position | SHL value |
|---|---|
| [5-8] | (SHL -6.5) |
| [15-18] | (SHL -5.5) |
| [26-29] | (SHL -4.5) |
| [37-40] | (SHL -3.5) |
| [47-50] | (SHL -2.5) |
| [57-60] | (SHL -1.5) |

**Major-groove bending sites**

| DNA position | SHL value |
|---|---|
| [10-14] | (SHL -6) |
| [21-23] | (SHL -5) |
| [32-34] | (SHL -4) |
| [42-45] | (SHL -3) |
| [52-55] | (SHL -2) |
| [62-65] | (SHL -1) |

Figure S1. Locations of the minor- and major-groove bending sites (GBS) in nucleosomal DNA. (A) The crystal structure of the 1KX5 nucleosome core particle (NCP) with 147-bp long DNA (Davey et al. 2002) shown schematically: the DNA fragment is divided into two halves, separated by the dyad (black ball and arrow). The base-pair centers in the 'ventral' half are represented by large balls, and the sugar-phosphate backbone is shown by a yellow ribbon. For the 'dorsal' half of the nucleosome, the base-pair centers are connected by sticks. Minor- and major-groove bending sites are shown in blue and red, respectively. These sites are named by their superhelical locations (SHL). (B) The exact locations of minor- and major-GBS in the 'ventral' half of the nucleosomal DNA fragment are shown. The sites on the 'dorsal' half are symmetrical to their counterparts on the 'ventral' half with respect to the dyad (Cui and Zhurkin 2010, Supplemental Table S1). The minor-groove bending sites at SHL ±0.5 (in grey) are not included for analysis because DNA patterns are out of phase at these locations (Satchwell et al. 1986).

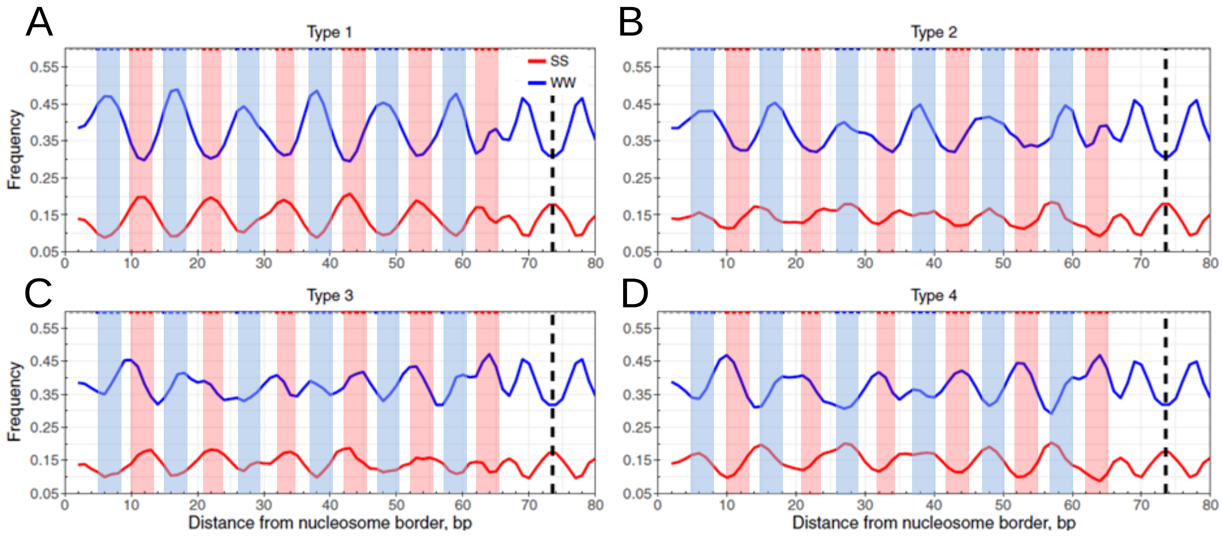

Figure S2. Four sequence patterns of yeast nucleosomal DNA mapped by a chemical method. The dyad positions of nucleosomes were published previously (Brogaard et al. 2010). Notations are the same as in Figure 1.

.

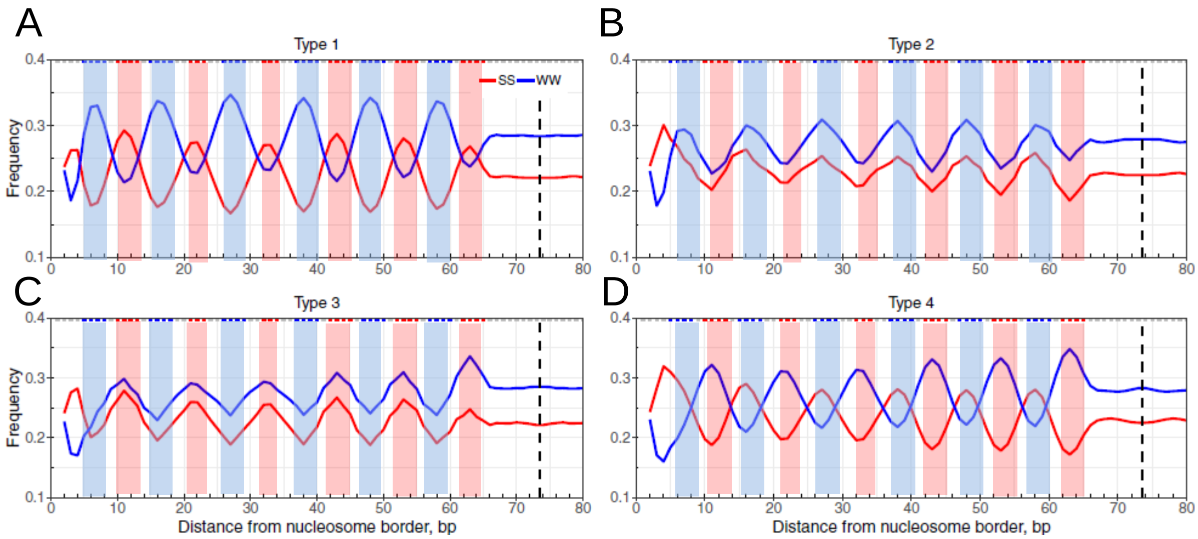

Figure S3. Four sequence patterns of fruit fly nucleosomal DNA mapped by MNase-Seq. The nucleosomal DNA fragments were sequenced by paired-end sequencing and the data were published previously (Chereji et al. 2015). Notations are the same as in Figure 1.

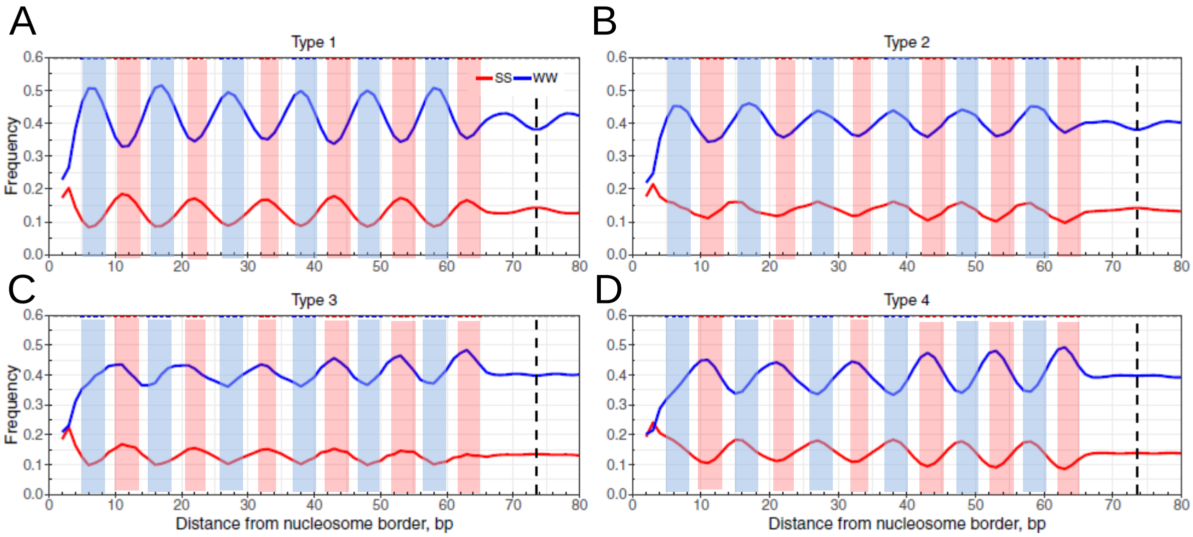

Figure S4. Four sequence patterns of nematode embryo nucleosomal DNA mapped by MNase-Seq. The nucleosomal DNA fragments were sequenced by paired-end sequencing and the data were published previously (Tabuchi et al. 2015). Notations are the same as in Figure 1.

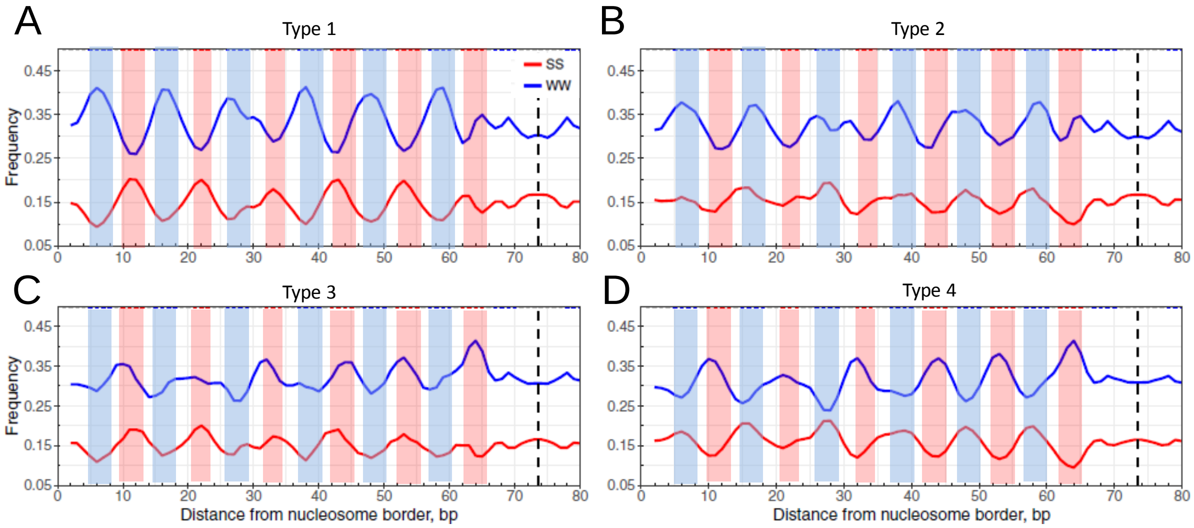

Figure S5. Four sequence patterns of mouse mESC nucleosomal DNA mapped by a chemical method. The dyad positions of nucleosomes were published previously (Voong et al. 2016). Notations are the same as in Figure 1.

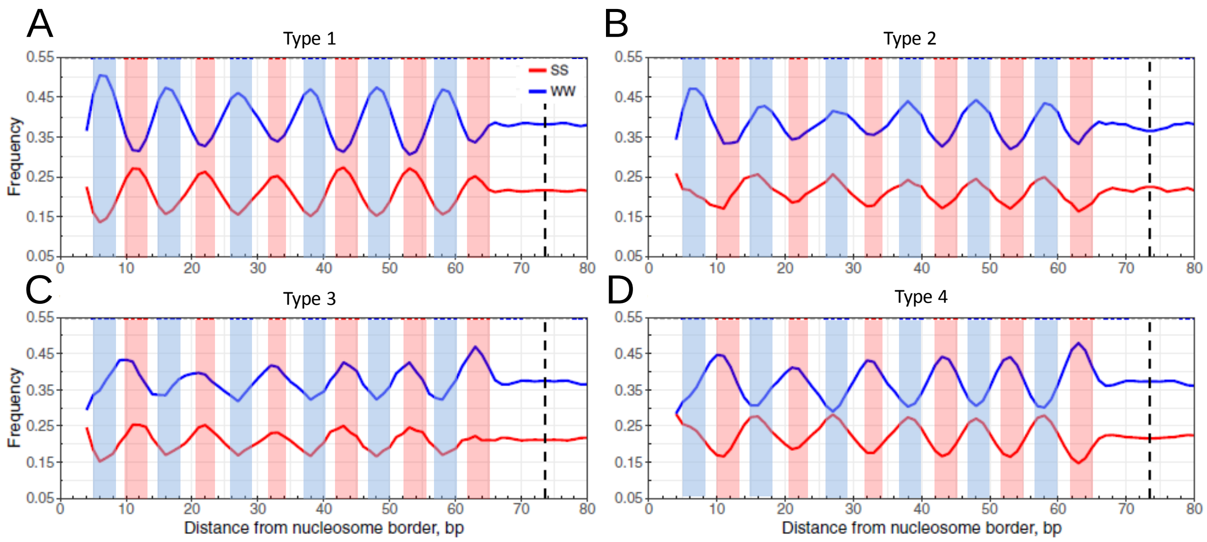

Figure S6. Four sequence patterns of mouse mESC nucleosomal DNA mapped by MNase-Seq. The nucleosomal DNA fragments were sequenced by paired-end sequencing and the data were published previously (Voong et al. 2016). Notations are the same as in Figure 1.

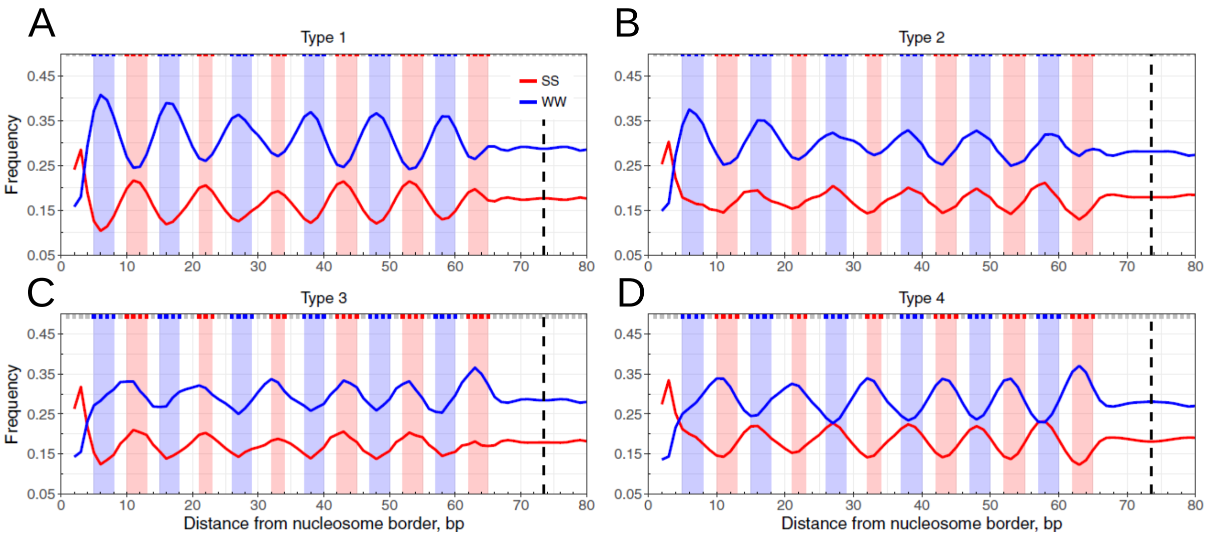

Figure S7. Four sequence patterns of human lymphoblastoid nucleosomal DNA mapped by MNase-Seq. The nucleosomal DNA fragments were sequenced by paired-end sequencing and the data were published previously (Gaffney et al. 2012). Notations are the same as in Figure 1.

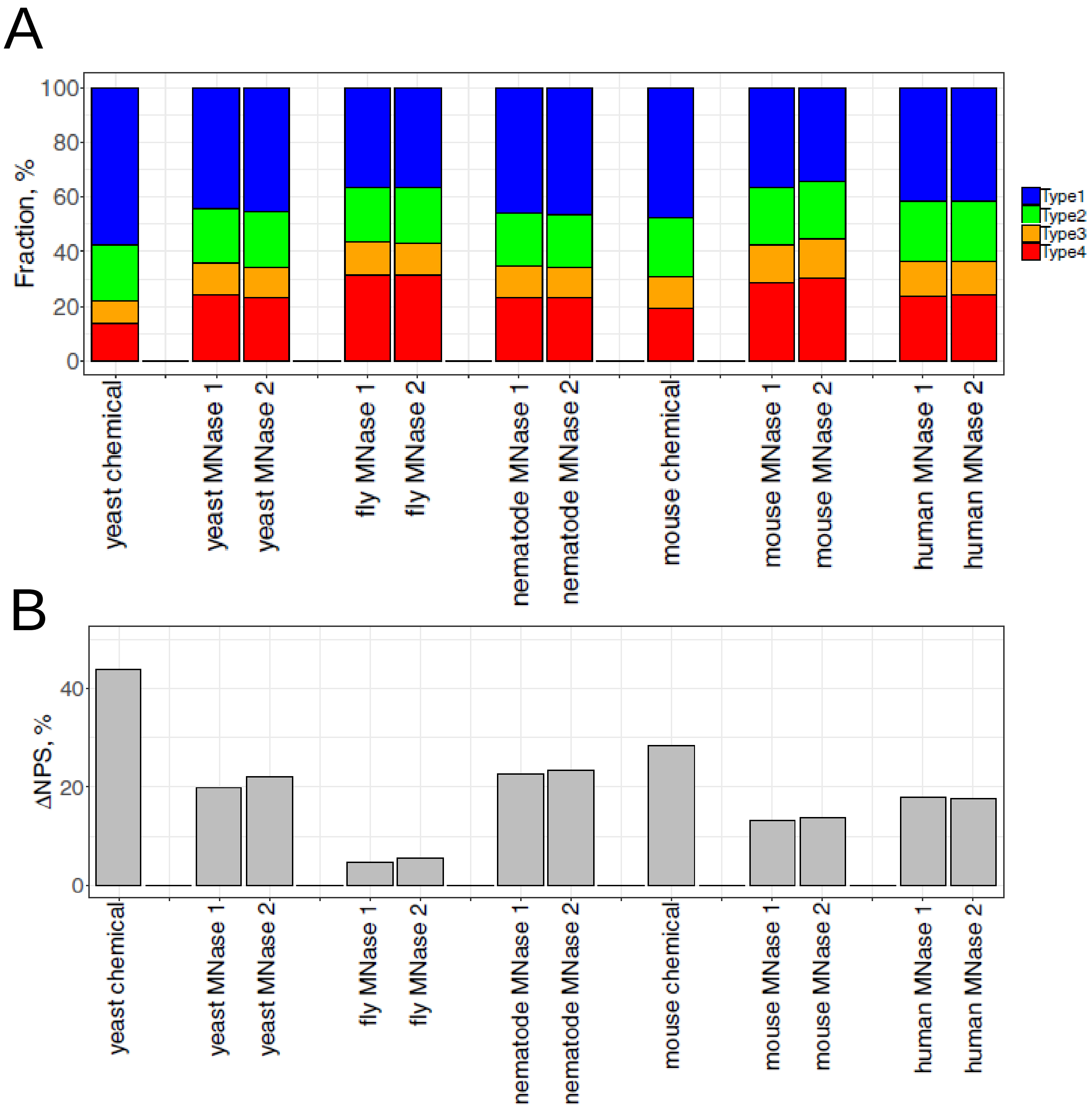

Figure S8. Comparison of the fractions of nucleosomal DNA patterns in eukaryotes. (A) Fractions of four sequence patterns in nucleosomal DNA from yeast, nematode, fruit flies, mice and humans. For chemical mapping data, the "unique" maps of nuleosomal dyad positions were taken from literature (Brogaard et al. 2010, Voong et al. 2016) and the corresponding 147-bp NCP fragments were used in this study. For the paired-end MNase mapping data, the two biological replicates or relevant datasets were taken from literature and 147-bp NCP fragments in these datasets were used for analysis. The fractions of sequence patterns were calculated for each dataset (Table S4). (B) ΔNPS value of each dataset. The ΔNPS is calculated as the difference between Type 1 nucleosomes (%) and Type 4 nucleosomes (%).

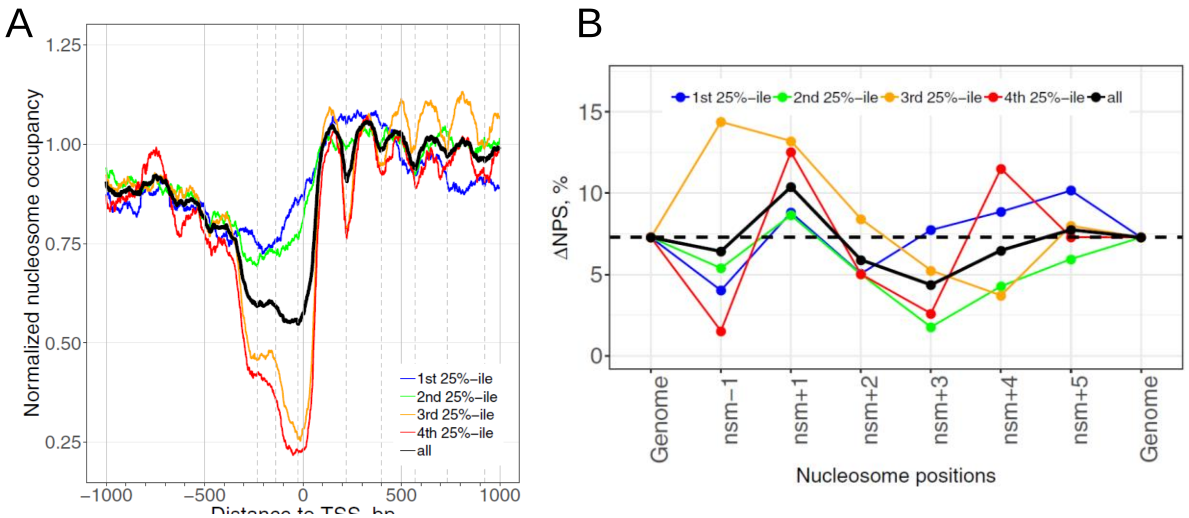

Figure S9. Nucleosome occupancy and ΔNPS values of fruit fly S2 cells. (A) Nucleosome occupancy profiles around TSS of fly. Nucleosomes mapped by MNase-Seq were taken from literature (Fuda and Lis 2015). Nucleosome occupancy signals ±1kb of verified TSSs are separated into quartiles by transcriptional frequencies based on RNA-seq data (Table S5). Notations are the same as Figure 2. (B) Nucleosome ΔNPS values in genes separated into quartiles by transcriptional frequencies. The ΔNPS values of all genes are shown in black. The genomic ΔNPS values are denoted by dashed lines.

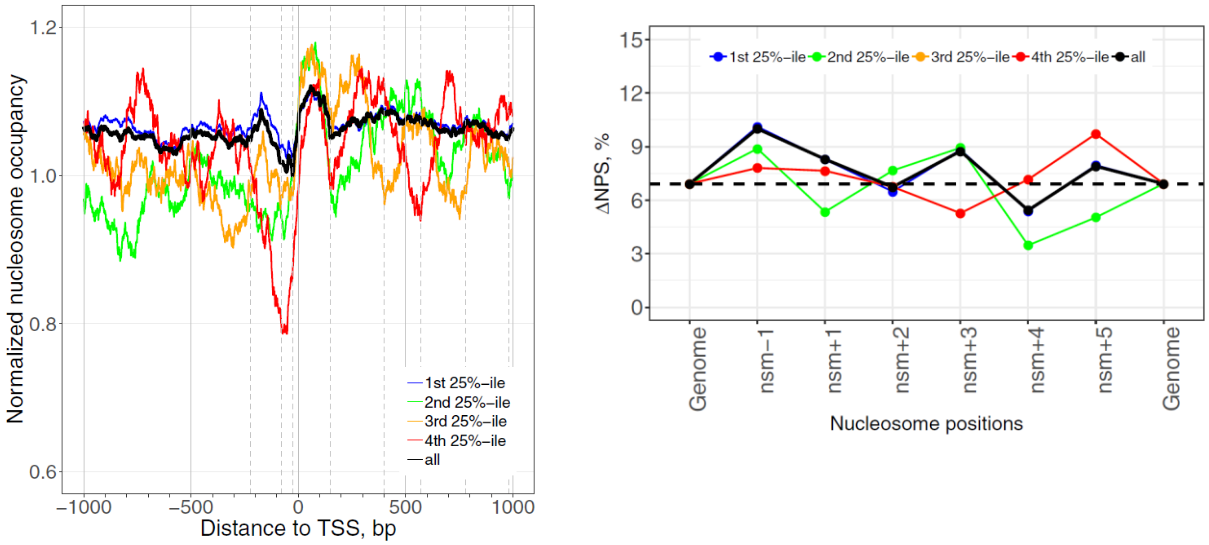

Figure S10. Nucleosome occupancy and ΔNPS values of nematode sperms. (A) Nucleosome occupancy profiles around TSS of fly. Nucleosomes mapped by MNase-Seq were taken from literature (Tabuchi et al. 2018). Nucleosome occupancy signals ±1kb of verified TSSs are separated into quartiles by transcriptional frequencies based on RNA-seq data (Table S5). Notations are the same as Figure 2. (B) Nucleosome ΔNPS values in genes separated into quartiles by transcriptional frequencies. The ΔNPS values of all genes are shown in black. The genomic ΔNPS values are denoted by dashed lines.

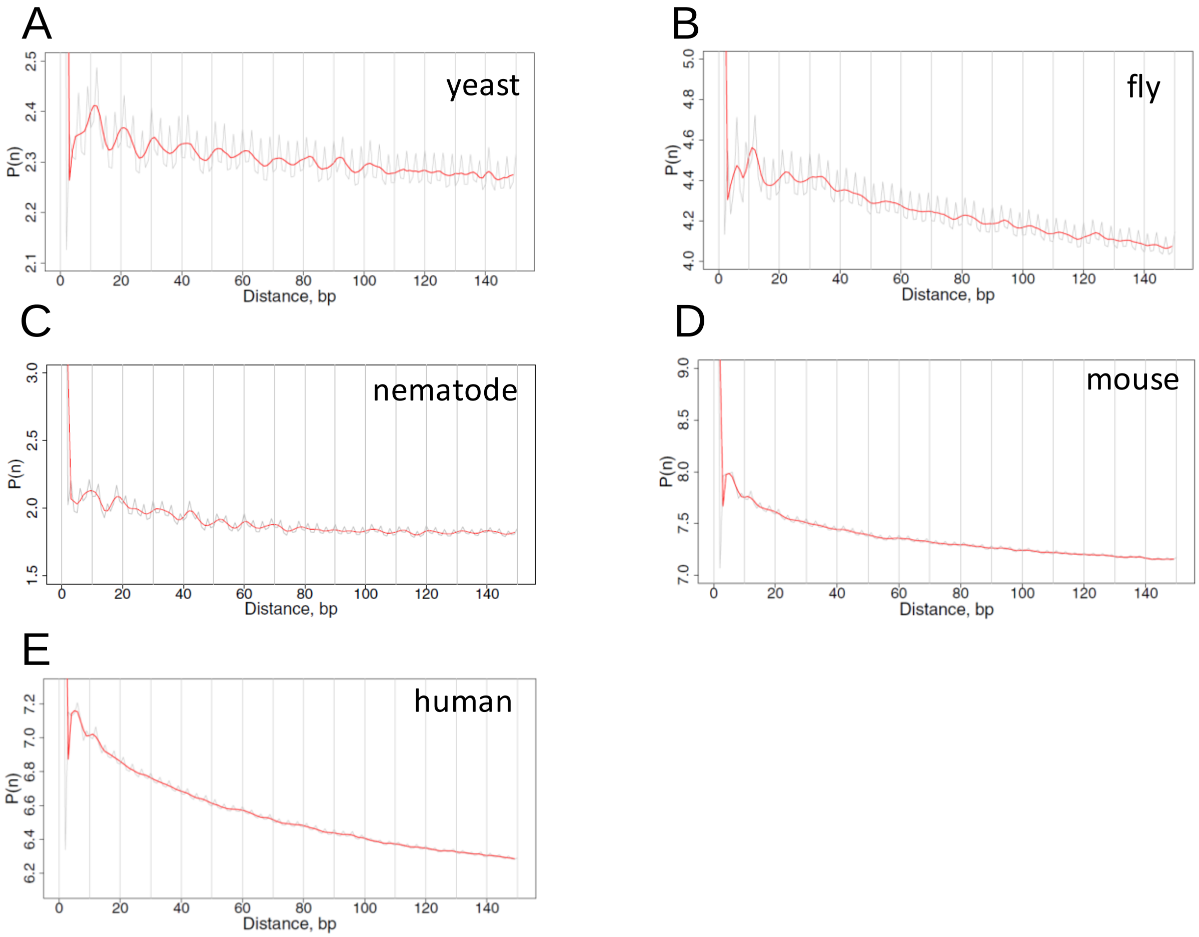

Figure S11. Distance auto-correlation function profiles for SS dinucleotides in yeast (A), fly (B), nematode (C), mouse (D) and human (E) DNA. Genomic fragments [-500 bp, +1000 bp] relative to verified TSSs (position 0) were used for analysis. Both raw (in gray) and 3-bp running average (in red) values were plotted. The distance auto-correlation function follows what was published before (Cui et al. 2012).

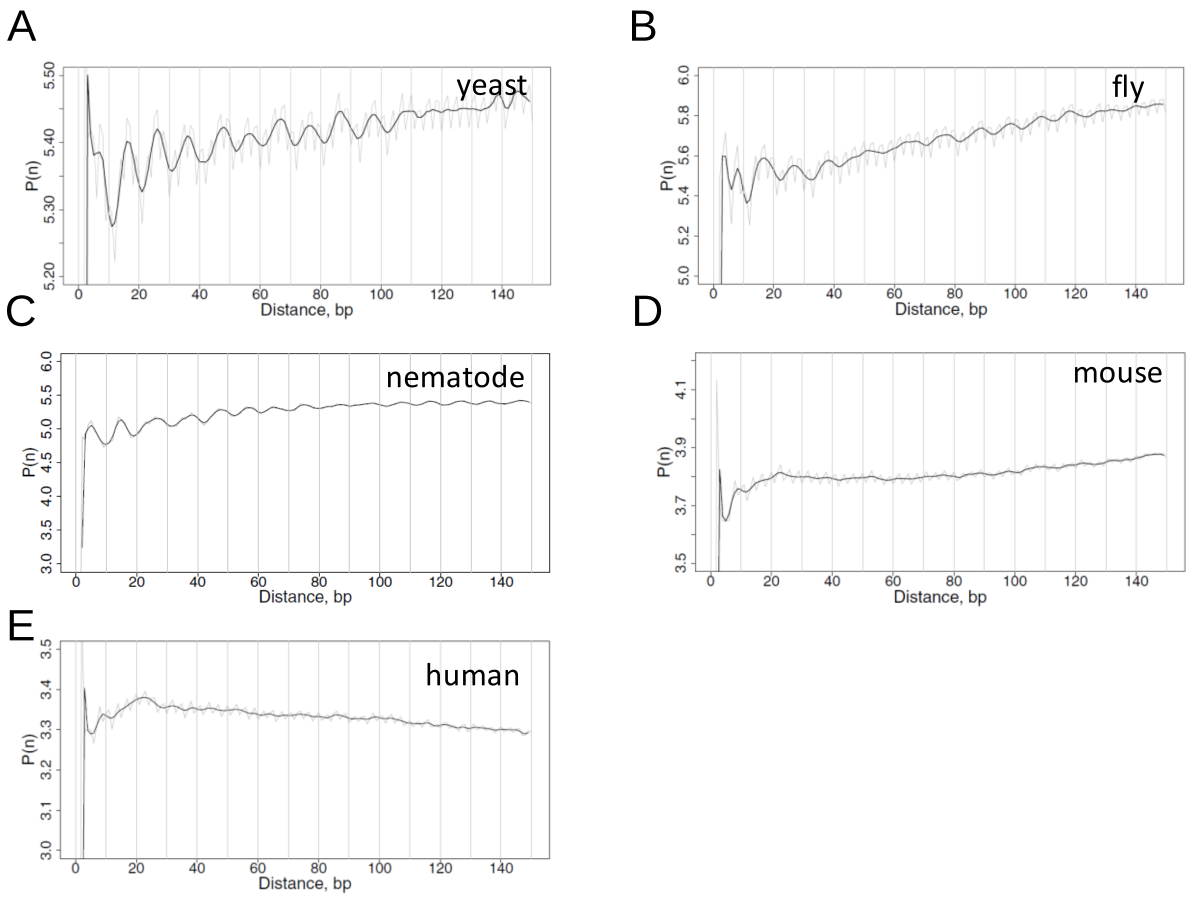

Figure S12. Distance cross-correlation function profiles for WW and SS dinucleotides in yeast (A), fly (B), nematode (C), mouse (D) and human (E) DNA. Genomic fragments [-500 bp, +1000 bp] relative to verified TSSs (position 0) were used for analysis. Both raw (in gray) and 3-bp running average (in black) values were plotted. The distance cross-correlation function follows what was published before (Cui et al. 2012).

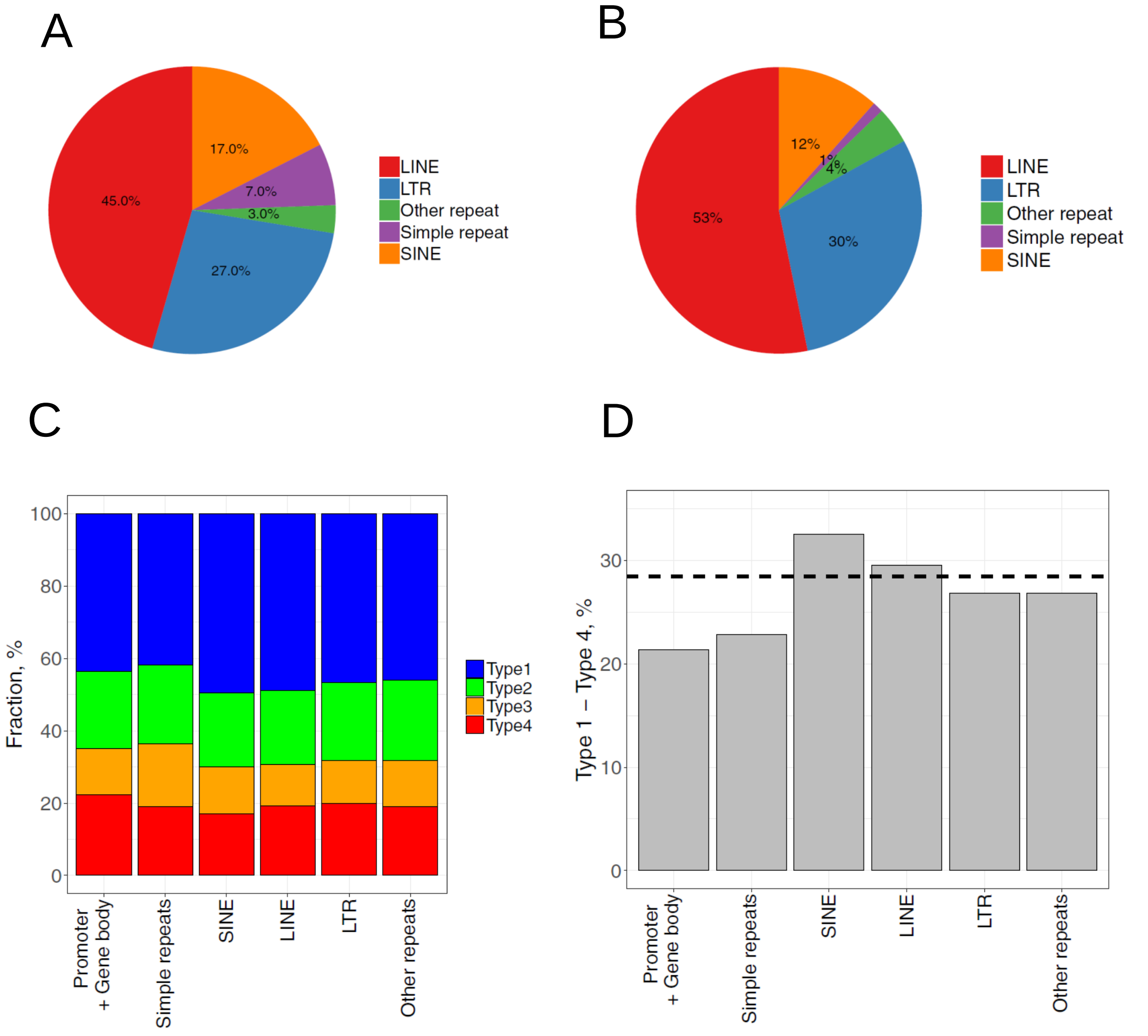

Figure S13. Various nucleosomal DNA sequence patterns in mouse transposable elements (TEs). (A) Fractions of mouse TE families. The fractions of TE families in the mouse genome were taken from literature (http://www.repeatmasker.org/species/mm.html). (B) Fractions of 147-bp mouse nucleosomes residing in TEs grouped by their families. (C) Fractions of 4 types of nucleosomes in TEs grouped by families. (D) Nucleosome ΔNPS values in genic and repetitive DNA regions. The genomic ΔNPS value is indicated by dashed lines.

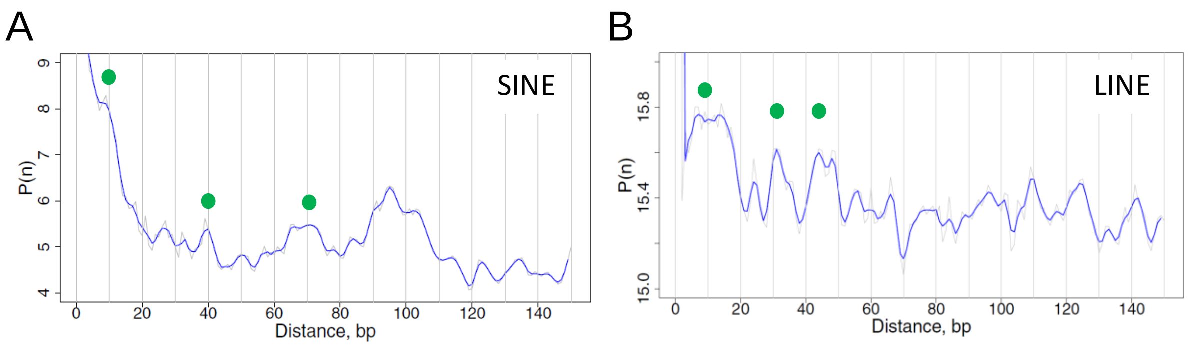

Figure S14. Distance auto-correlation functions for WW dinucleotides in human transposable elements. SINE (A) and LINE (B) elements from human genome (hg18) were used for analysis. Both raw (gray) and 3-bp running averages (blue) of the function values were profiled. Green dots indicate that neighboring WW dinucleotides are separated by a multiple of ~10 bp.

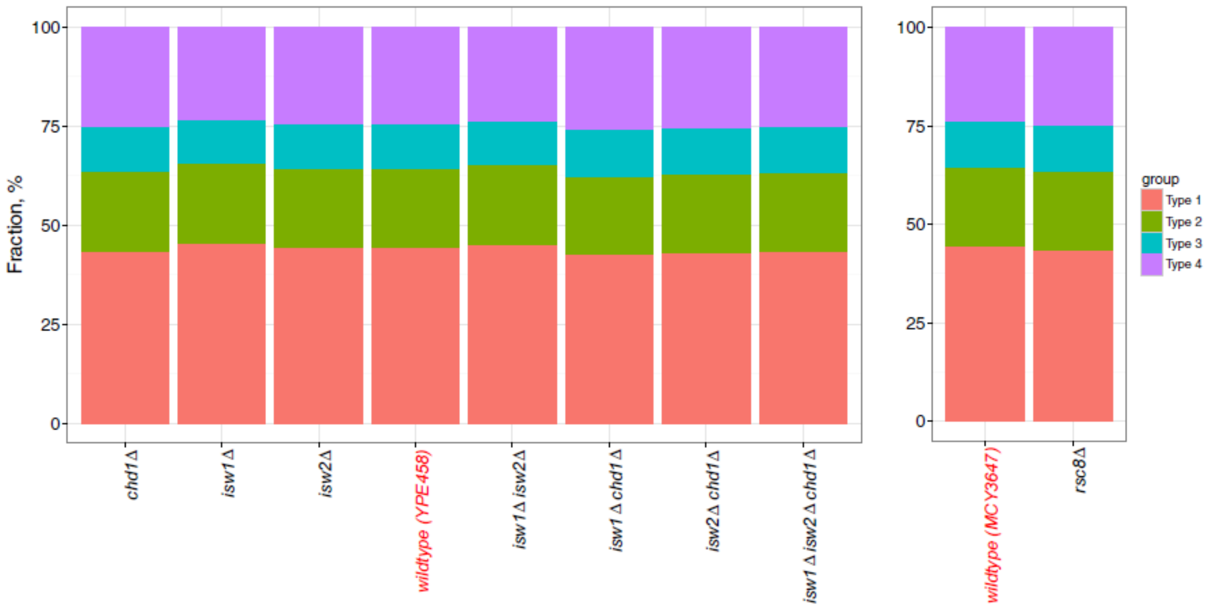

Figure S15. Fractions of 4 types of nucleosomal DNA in yeast wildtype strains and mutants. Notations are the same as in Figure S8A.

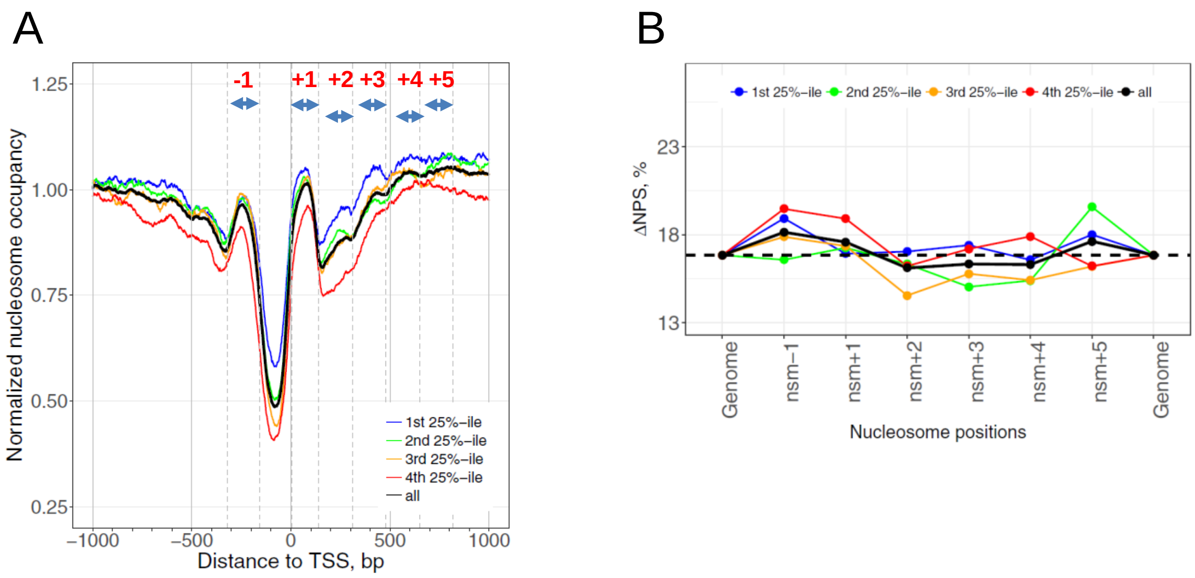

Figure S16. Nucleosome occupancy profiles and ΔNPS values around TSS in a yeast mutant strain (*isw1Δ chd1Δ* double mutant) (Ocampo et al. 2016). (A) Nucleosome occupancy signals ±1 kb of verified TSSs. For the sake of comparison, the ranges of nucleosomes are the same for both the wildtype and mutant strains (Table S6). Other notations follow Figure 3. (B) Nucleosome ΔNPS values in genes separated into quartiles by transcriptional frequencies. The ΔNPS values of all genes are shown in black. The genomic ΔNPS values are denoted by dashed lines.

**Supplementary References**

Brogaard K, Xi L, Wang JP, Widom J. 2012. A map of nucleosome positions in yeast at base-pair resolution. *Nature* **486**: 496-501.

Chereji RV, Kan TW, Grudniewska MK, Romashchenko AV, Berezikov E, Zhimulev IF, Guryev V, Morozov AV, Moshkin YM. 2016. Genome-wide profiling of nucleosome sensitivity and chromatin accessibility in Drosophila melanogaster. *Nucleic Acids Res* **44**: 1036-1051.

Cui F, Zhurkin VB. 2010. Structure-based analysis of DNA sequence patterns guiding nucleosome positioning in vitro. *J Biomol Struct Dyn* **27**: 821-841.

Cui F, Cole HA, Clark DJ, Zhurkin VB. 2012. Transcriptional activation of yeast genes disrupt intragenic nucleosome phasing. *Nucleic Acids Res* **40**: 10753-10764.

Davey CA, Sargent DF, Luger K, Maeder AW, Richmond TJ. 2002. Solvent mediated interactions in the structure of the nucleosome core particle at 1.9Å resolution. *J Mol Biol* **391**: 1097-1113.

Fuda NJ, Guertin MJ, Sharma S, Danko CG, Martins AL, Siepel A, Lis JT. 2015. GAGA factor maintains nucleosome-free regions and has a role in RNA polymerase II recruitment to promoters. *PLoS Genet.* **11**: e1005108.

Gaffney DJ, McVicker G, Pai AA, Fondufe-Mittendorf YN, Lewellen N, Michelini K, Widom J, Gilad Y, Pritchard JK. 2012. Controls of nucleosome positioning in the human genome. *PLoS Genet* **8**: e1003036.

Ganguli D, Chereji RV, Iben JR, Cole HA, Clark DJ. 2014. RSC-dependent constructive and destructive interference between opposing arrays of phased nucleosomes in yeast. *Genome Res* **24**: 1637-1649.

Ocampo J, Chereji RV, Eriksson PR, Clark DJ. 2016. The ISW1 and CHD1 ATP-dependent chromatin remodelers compete to set nucleosome spacing in vivo. *Nucleic Acids Res* **44**: 4625-4635.

Satchwell SC, Drew HR, Travers AA. 1986. Sequence periodicities in chicken nucleosome core DNA. *J Mol Biol* **191**: 659-675.

Tabuchi TM, Rechtsteiner A, Jeffers TE, Egelhofer TA, Murphy CT, Strome S. 2018. Caenorhabditis elegans sperm carry a histone-based epigenetic memory of both spermatogenesis and oogenesis. *Nat Commun* **9**: 4310.

Voong LN, Xi L, Sebeson AC, Xiong B, Wang JP, Wang X. 2016. Insights into nucleosome organization in mouse embryonic stem cells through chemical mapping. *Cell* **167**: 1555-1570.
